# Structured Connectivity Across Multiple *Drosophila* Mushroom Bodies

**DOI:** 10.64898/2026.09.06.749654

**Authors:** M.W. Pleijzier, P. Schlegel, L. Serratosa Capdevila, Cambridge Connectomics Group, FlyEM Project Team, K.D. Longden, G.S.X.E. Jefferis

## Abstract

Learning and memory depend on plasticity within neural circuits, but the extent to which the underlying architecture of memory networks is stereotyped or structured remains unclear. Random connectivity may maximise stimulus discrimination, whereas structured connectivity could preferentially route behaviourally relevant information. Here, we compare five adult *Drosophila melanogaster* Mushroom Body (MB) hemispheres across three connectomes and establish a unified cross-connectome framework for uniglomerular projection neurons (uPNs), Kenyon Cells (KCs), Mushroom Body Output Neurons (MBONs) and Lateral Horn Centrifugal Neurons (LHCENTs). Despite variation in KC abundance and fine-scale connectivity, we identify reproducible organisation across the uPN→KC and KC→MBON pathways. A canonical group of uPNs encoding odours associated with food was consistently identified across hemispheres and has non-random connectivity onto KCs. This bias is preferentially routed through KCs towards approach-promoting MBONs. Sensory input also remains stereotyped across sister MBONs from the same hemisphere despite variation in their individual KC partners. Finally, recurrent LHCENT feedback (predicted-inhibitory) preferentially targets KCs receiving these food-associated inputs. Together, these findings show that a memory network can accommodate local variability while preserving structured pathways that prioritise ethologically relevant sensory information.

## Introduction

Synaptic modification and circuit remodelling underlie learning and memory (Uytiepo *et al*., 2025), but circuit-level variation is difficult to resolve without complete synapse-resolution data. For example, David Marr assumed random connectivity between input channels and downstream interneurons in the absence of high-quality connectivity data from the cerebellar cortex, noting that “it is regrettable that no data exist to suggest a better model” (Marr, 1969). Random connectivity, defined by the absence of preferential targeting between neuronal layers, has been asserted for memory systems including the cerebellum, hippocampus, olfactory bulb and insect Mushroom Body (MB; Stevens, 2015). Sparse random connectivity can increase representational dimensionality and stimulus discrimination (Litwin-Kumar *et al*., 2017), making its prevalence in biological memory networks a central question.

The insect MB supports associative learning and memory and is well-defined in *Drosophila melanogaster*, making this system ideal for exploring patterns of neural connectivity (Figure 1A). Each antennal lobe (AL) comprises ∼50 glomeruli, each encoding a characteristic set of odours (Couto *et al*., 2005; Hallem and Carlson, 2006). ∼120 projection neurons (PNs) relay olfactory information to ∼2,000 Kenyon Cells (KCs) in the Calyx (Figure 1A). KC axons traverse the MB lobes and contact MB output neurons (MBONs). Three major morphological KC cell types are defined by the MB lobes (α, β, α′, β′ and γ) they occupy: KCαβ, KCα′β′ and KCγ. Compartmentalised dopaminergic input regulates KC→MBON plasticity and supports learned behaviours (Cognigni *et al*., 2018). Recent work showed that MBONs then connect to Lateral Horn Centrifugal Neurons (LHCENTs) downstream of the MB (Bates *et al*., 2020).

**Figure 1:**
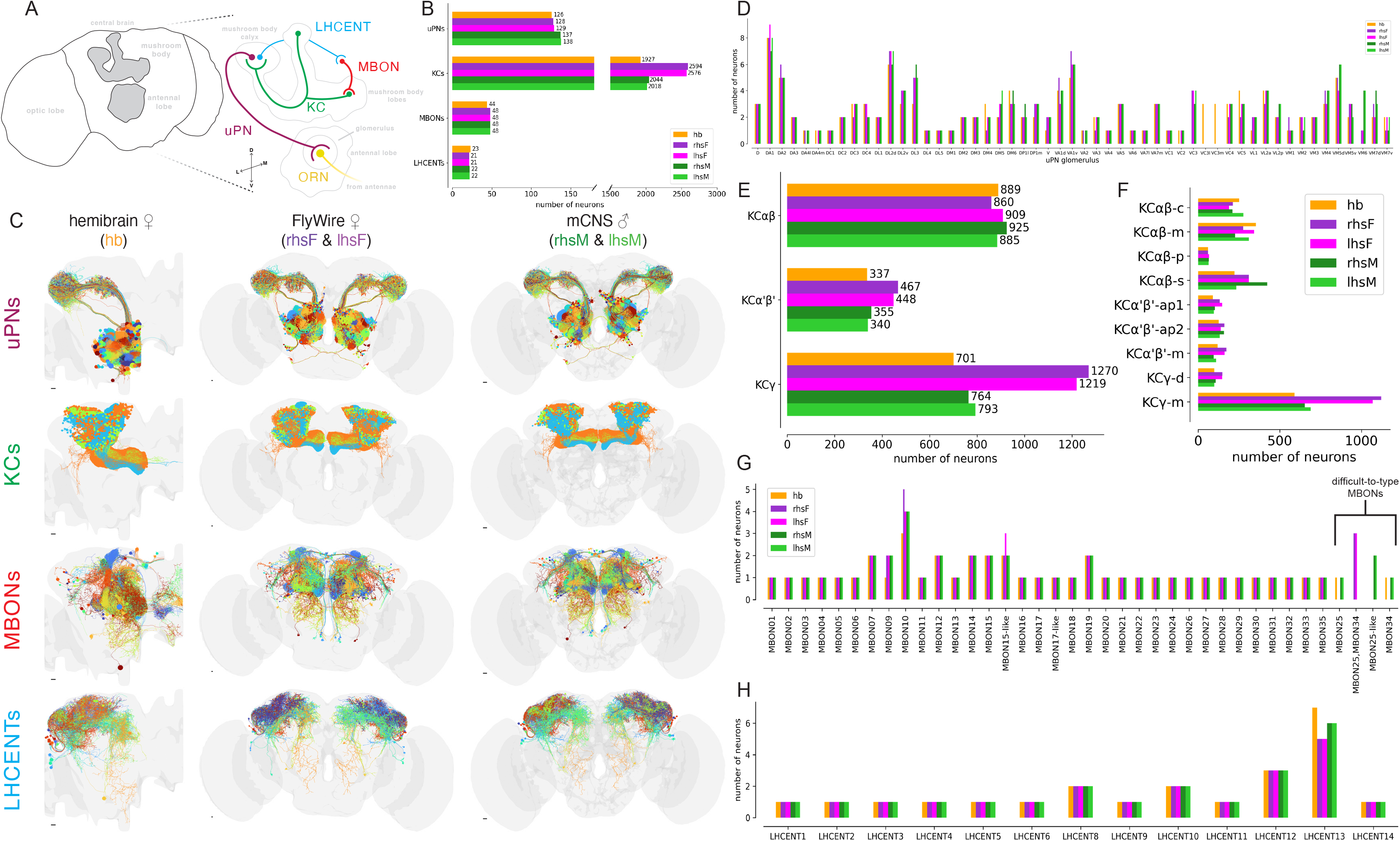
Cross-hemisphere stereotypy of the major cell types involved in olfactory-mnemonic processing: **(A)** Illustrative olfactory-mnemonic circuit in *Drosophila melanogaster*: Olfactory Receptor Neurons (ORNs) connect onto projection neurons (PNs) in the antennal lobe. We focus on uniglomerular projection neurons (uPNs) that relay olfactory information to the Mushroom Body (MB) Calyx, where they synapse onto the intrinsic neurons of the MB, Kenyon Cells (KCs). KCs send axons into the MB lobes where they synapse onto the dendrites of Mushroom Body Output Neurons (MBONs). MBONs then send axons outside of the MB into the superior protocerebral areas (SLP/SIP/SMP), where they connect onto the dendrites of Lateral Horn Centrifugal Neurons (LHCENTs). Recurrent LHCENTs then send axons back into the Calyx, where they connect onto KCs. **(B)** Number of uPNs, KCs, MBONs and LHCENTs across hemispheres. **(C)** Morphologies of each of the major olfactory-mnemonic cell types in each dataset. Colours are assigned independently across the major cell types (for KCs, blue = KCαβ, green = KCα′β′, orange = KCγ) and the scale bars represent 10 μm. Number of neurons in each **(D)** uPN cell type, **(E)** broad KC type, **(F)** KC subtype, **(G)** MBON type and **(H)** LHCENT type, across the five hemispheres. For MBON types, the difficult-to-type neurons are indicated with the black brackets, and morphology plots of these neurons are found in Supplementary Figure 1.1G.

Single-cell labelling studies have demonstrated that olfactory PN axons have stereotyped, zonal arborisation patterns at discrete locations in the adult Calyx (Lin *et al*., 2007; Jefferis *et al*., 2007; Marin *et al*., 2002; Wong *et al*., 2002). Recent advances in adult *Drosophila* EM-based connectomics support this description of non-random connectivity: a partial reconstruction of the full adult fly brain (FAFB) EM volume revealed that overlapping populations of KCs are targeted by PNs conveying information from glomeruli encoding food-related odours (Zheng *et al*., 2022); meanwhile, visual inputs to the MBs target a non-random subset of KCs in the female hemibrain and FlyWire datasets (Ganguly *et al*., 2024). However, a study tracing connections from KCs back to PNs found no structured organisation, suggesting random PN→KC connectivity (Caron *et al*., 2013). This description of random connectivity was supported by a study using *Orco^−/−^*flies, showing that PN→KC connectivity developed independently of olfactory experience (Hayashi *et al*., 2022). An analysis of a whole-brain larval *D. melanogaster* connectome also demonstrated that most KCs integrate a random combination of PN inputs (Eichler *et al*., 2017). However, the MB is extensively remodelled during metamorphosis (Truman *et al*., 2023), leaving the question of whether random PN→KC connectivity persists into adulthood unresolved. Finally, the initial analysis of the FAFB dataset (Zheng *et al*., 2022) may have been biased by sampling only ∼52% (1,356) of the KCs subsequently identified in the fully proofread dataset (Schlegel *et al*., 2024; Dorkenwald *et al*., 2024).

Here, we resolve these contradictory findings through comprehensive analysis of five fully reconstructed MB circuits across three synapse-resolution connectomes and five hemispheres: the female hemibrain (labelled as ‘hb’, Li *et al*., 2020; Scheffer *et al*., 2020), the bilateral female FlyWire (labelled as ‘lhsF’ and ‘rhsF’, Dorkenwald *et al*., 2024; Schlegel *et al*., 2024) and the bilateral male central nervous system (mCNS, labelled as ‘lhsM’ and ‘rhsM’, Berg *et al*., 2026). We first establish a unified cell typing of MB-associated neurons, enabling fair comparisons of cell-type counts, dataset-specific differences and KC-input organisation. We show that while many adult PN→KC connections are randomly organised, a subset of PN glomeruli that transmit food-related odour information is reliably non-random across hemispheres, individuals and sexes. We then show that this structured input connectivity to the MB is preserved in MBON inputs, the primary sites of memory formation, and identify a conserved bifurcation in effective PN→KC→MBON routing. Finally, we characterise an inhibitory feedback pathway from MBONs→LHCENTs→KCs that complements the structured input from food-odour-encoding PNs onto KCs in the Calyx. Using comparative connectomics, we comprehensively show that stereotyped, non-random patterns of connectivity support the processing of ethologically relevant food odours in a memory network with otherwise randomly organised input connections.

## Results

### A unified consensus and numerical stereotypy of MB cell types

To measure how cell-type connectivity varies across hemispheres and datasets, we first established consensus identities and cell counts (i.e. numerical stereotypy) of four major olfactory-mnemonic populations: uniglomerular olfactory projection neurons (uPNs), KCs, MBONs and LHCENTs (Figure 1). To do this, we integrated hemibrain (hb), FlyWire (lhsF and rhsF) and mCNS cell type catalogues (lhsM and rhsM).

uPNs were typed by the AL glomerulus they innervate, and we accounted for missing or inconsistent annotations, such as V, MV6 and VC3. Total uPN counts varied only by 5% (126–133 neurons per hemisphere; Figure 1B). The number of neurons within each uPN type was also consistent, varying ±1 between hemispheres (Figure 1D). uPN numerical stereotypy varied with the glomerulus birth order, as suggested in previous work (Supplementary Figure 1.1A–F; Schlegel *et al*., 2021). In contrast, KC cell type abundance varied substantially between datasets. There were 26–35% more cells per hemisphere in FlyWire, compared with hemibrain and mCNS (Figure 1B). Major KC cell types (KCαβ, KCα′β′ and KCγ) can be further subdivided into morphological subtypes (Li *et al*., 2020): KCαβ-c, KCαβ-m, KCαβ-p, KCα′β′-ap1, KCα′β′-ap2, KCα′β′-m, KCγ-m, KCγ-d, KCγ-s1/2/3/4 and KCγ-t. The higher KC count primarily reflected 54-81% more KCγ neurons in FlyWire, largely owing to expansion of KCγ-m (Figure 1E–F). Additionally, KCα′β′ were slightly increased in FlyWire, whereas KCαβ were stable (860–917). As the mCNS closely matched the female hemibrain, FlyWire appears to be a biological outlier (in terms of KC number) rather than a new baseline in KC abundance. Singleton subtypes KCγ-s1 and KCγ-s2 (not shown) were recovered across datasets, but KCγ-s3, KCγ-s4 and KCγ-t were only found in hemibrain, suggesting hemibrain oversegmentation artefacts and illustrating the value of cross-connectome typing.

Most typical and atypical MBONs were recovered across hemispheres and occurred as one neuron per type per hemisphere, consistent with light-level studies (Figure 1G; Aso *et al*., 2014a). Exceptions to this included variable counts for MBON09, MBON10 and MBON15-like: there was only one hemibrain MBON09, compared to two cells per hemisphere in FlyWire and mCNS; MBON10 varied by ±2 neurons per hemisphere; and there were three MBON15-like neurons in the left FlyWire hemisphere, but two in the other hemispheres (Figure 1G). A bilateral, mCNS-specific, MBON25-like was absent from the other datasets, but we did not interpret it as a male-specific cell type. Instead, MBON25, MBON25-like and MBON34 collectively form a group of MBONs with highly similar morphology and connectivity because of their projections in the Crepine, a neuropil that wraps around the MB horizontal lobe (Ito *et al*., 2014). These types are most reliably distinguished by their γ-lobe projections (Supplementary Figure 1.1G), as quantitative clustering alone could not separate them.

Most LHCENT types were also numerically stereotyped, usually with one neuron per type per hemisphere (Figure 1H). LHCENT8 and LHCENT10 had two neurons, LHCENT12 had three, and LHCENT13 varied modestly across datasets. These neurons extend dendrites through the SLP, SIP and SMP and project to the lateral horn; a subset also returns to the Calyx (Figure 7A; Bates *et al*., 2020). Across uPNs, KCs, MBONs and LHCENTs, we found no sex-associated differences in within-type neuron number, and the remaining variation likely reflects biological and/or minor technical effects. Together, these data demonstrate a high degree of cell-type stereotypy across hemispheres and datasets, with noted instances of variability, such as KCγ types. These findings provide the foundation for the comparative connectivity analyses that follow.

### Connectome-specific differences in KC input budgets

Having systematically identified MB cell types across the datasets, we next quantified and compared uPN→KC connectivity across hemispheres, specifically examining how the greater number of KCs in FlyWire affected this connectivity pathway (Figure 1B). Accordingly, we grouped uPNs by the AL glomerulus they innervate and summed their connections to KCs. Total uPN input showed distinct patterns across KC classes and hemispheres. KCγ received the greatest input overall, averaging ∼84,600 ± 11,800 synapses across the five hemispheres, followed by KCαβ with 65,700 ± 8,100 synapses (mean ± SD; Figure 2A). These totals varied relatively little across hemispheres (coefficients of variation (CVs) of 14% and 12%, respectively). By contrast, KCα′β′ received substantially less total input and showed greater cross-hemisphere variation, averaging 17,200 ± 5,600 synapses and ranging from 10,800 to 23,800 (CV = 33%). Normalising for the number of neurons in each KC cell type revealed a similarly consistent mean input per KCαβ (78.9 ± 8.3 synapses per neuron; Figure 2B). Mean input was lower and more variable for KCα′β′ (47.9 ± 20.5;) and higher for KCγ (106.7 ± 26.6), with CVs of 43% and 25%, respectively.

**Figure 2:**
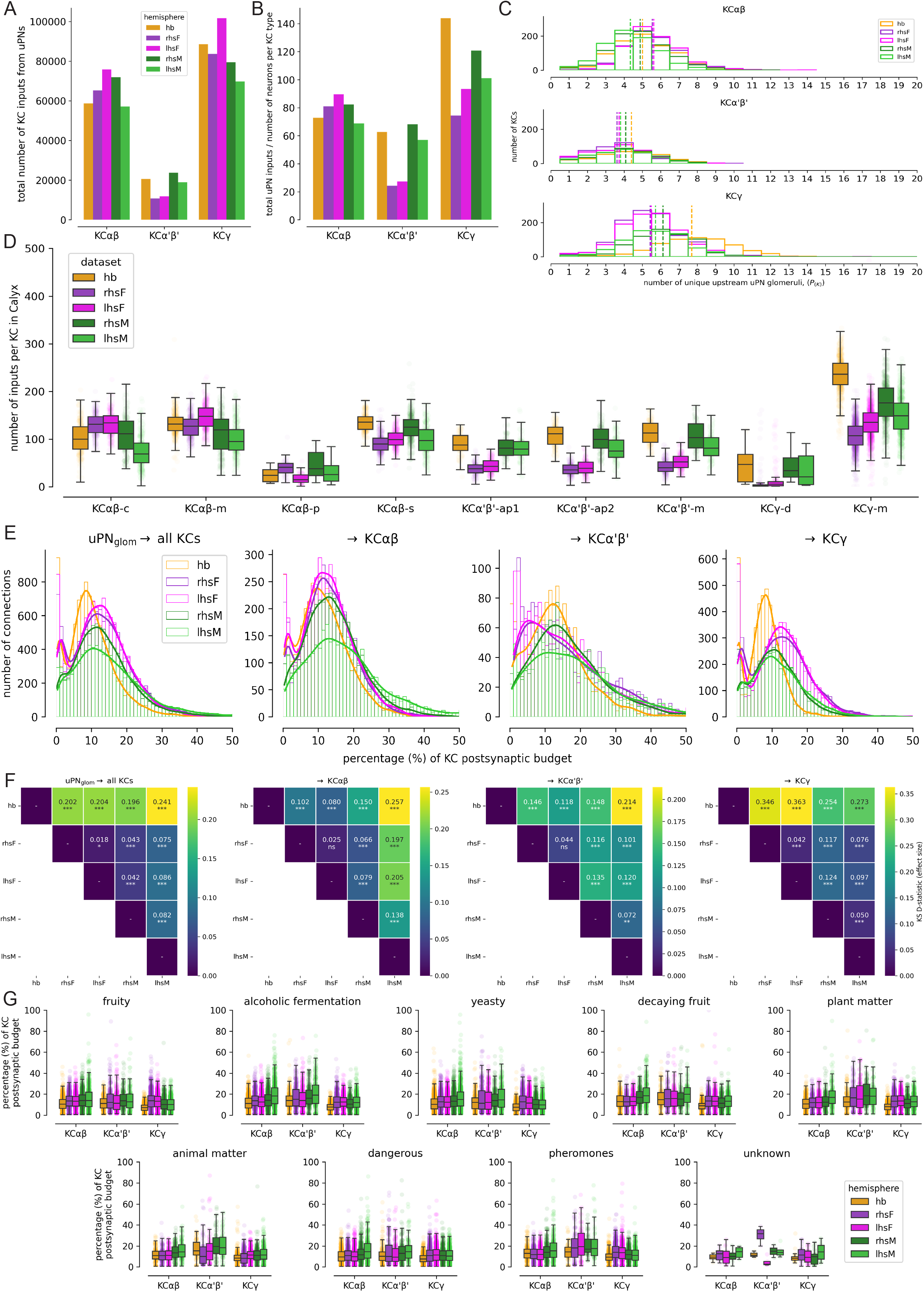
Variability in uPN glomerulus→KC connectivity across hemispheres: **(A)** Total number of inputs onto KCs from uPNs. **(B)** The average number of inputs per neuron for each broad KC type. This was calculated as the total number of uPN inputs per broad KC type divided by the total number of KC neurons in that type. **(C)** The number of unique upstream uPN glomeruli/types (i.e. odour channels) for each KC: for KCαβ (top row), KCα′β′ (middle row) and KCγ (bottom row). Dashed vertical lines represent the means of the distributions per hemisphere. **(D)** Distribution of the number of uPN synapse inputs onto KC subtypes. **(E)** Frequency distributions of the percentage of synaptic inputs from each uPN glomerulus onto individual KCs for all individual KCs (first panel), KCαβ (second panel), KCα′β′ (third panel) and KCγ (fourth panel) neurons. Bins are 1% of postsynaptic input in width. **(F)** *D*-statistics (effect sizes) from pairwise Kolmogorov-Smirnov (KS) tests on uPN→KC connectivity distributions presented in **(E)**. KS tests were performed with Benjamini-Hochberg false-discovery-rate (BH-FDR) correction (see Supplementary Figure 2.1 for the *p*-value matrices). Asterisks denote significance: *** if *p* < 0.001, ** if *p* < 0.01, * if *p* < 0.05, otherwise ‘ns’ for non-significant. **(G)** Percentage of KC postsynaptic budget by odour scene across hemispheres. For each hemisphere, uPN glomerulus→KC connectivity matrices were normalised by the total number of inputs received by each individual KC neuron in the Calyx (i.e. the fraction of inputs) and converted into percentages. uPN glomeruli were assigned to odour scenes using the “odour_scenes” table (see Methods), and each KC was assigned to its corresponding KC type. Boxplots show medians and interquartile ranges. Whiskers extend to the most extreme values within 1.5× the interquartile range, and the points behind the boxplots show the distribution of values.

To explore how these differences alter the number of unique odour input channels onto KCs, we compared the distributions of the number of unique glomeruli providing uPN synaptic input to each KC for the KCαβ, KCα′β′ and KCγ cell types (Figure 2C). The numbers of unique glomerular uPN inputs onto KCγ were more similar between the mCNS and FlyWire than between either dataset and hemibrain (pairwise Cohen’s *d* = 0.71–1.1; Figure 2C; Supplementary Figure 2.1C). To quantify how individual cell types contribute, we compared glomerular uPN inputs to individual KCs, normalised by the total number of input synapses each KC has in the Calyx (Figure 2E). For connections from uPN glomeruli to all individual KCs, the largest difference was between hemibrain and lhsM (Kolmogorov-Smirnov test, *D* = 0.241; Figure 2F), and across KCαβ, KCα′β′ and KCγ cell types, hemibrain KCs had the most divergent glomerular uPN-input distributions (Figure 2F).

To explore how these variations in MB input connectivity relate to the structural organisation of odour encoding in the ALs, we grouped glomeruli into “odour scenes” (e.g. “plant-matter”, “fruity”; Figure 2G), a grouping made possible by extensive prior work characterising the receptors expressed by olfactory receptor neurons (ORNs) that selectively innervate a glomerulus (see Methods for details). Within each odour scene and KC type, we compared the distributions of per-KC proportional uPN input across the five hemispheres. No detectable hemisphere effect was found in 13 of the 27 odour-scene × KC-type comparisons, whereas 14 showed a significant effect after correction for multiple comparisons (Kruskal-Wallis tests with Benjamini-Hochberg false discovery rate (BH-FDR) correction, *q* < 0.05; Supplementary Table 1). However, all detected effects were negligible (10 comparisons) or small (four comparisons), with no moderate or large effects. Therefore, although the percentage-input distributions for each KC type were not identical across odour scenes (Figure 2G), the magnitude of these differences was consistently limited across the five connectomes.

Together, these data demonstrate differences in fine-scale organisation of glomerular uPN inputs to KCs across the five hemispheric circuits, and show that mCNS and FlyWire had more similar uPN→KC connectivity to one another than either had to hemibrain (Figure 2E), despite differences in KC cell type numbers (Figure 1). They also demonstrate that when glomerular uPN input to KCs was grouped by odour scene, it varied little across KC types and datasets (Figure 2G). Across these analyses, we did not find evidence of sexual dimorphism in glomerular uPN inputs to KCs (Figure 2).

### Structured, non-random uPN→KC connectivity

Having compared the synaptic connectivity distributions between hemispheres, we investigated whether structured, non-random connectivity between uPNs and KCs in the adult Calyx was stereotyped across connectome hemispheres and individuals (Figure 3A). To detect circuit structure, we applied consensus cosine clustering, a method that quantified how frequently each pair of glomeruli co-clustered across all five hemispheres (see Methods for details; Monti *et al*., 2003; Strehl and Ghosh, 2002). This method yielded a set of four glomerular groups, which we refer to as “canonical” clusters (Supplementary Figure 3.1). One of these canonical clusters, the G2 cluster, frequently co-associated across hemispheres and comprised DM5, DP1l, VM3, VM2, VA4, VA2, DP1m, DM4, DM3, DM1, DM2, VC2, DL2v and DM6 (Figure 3B; Supplementary Figure 3.1B). The three other clusters (G1, G3 and G4) had less cohesive partitions (Figure 3B; Supplementary Figure 3.1).

**Figure 3:**
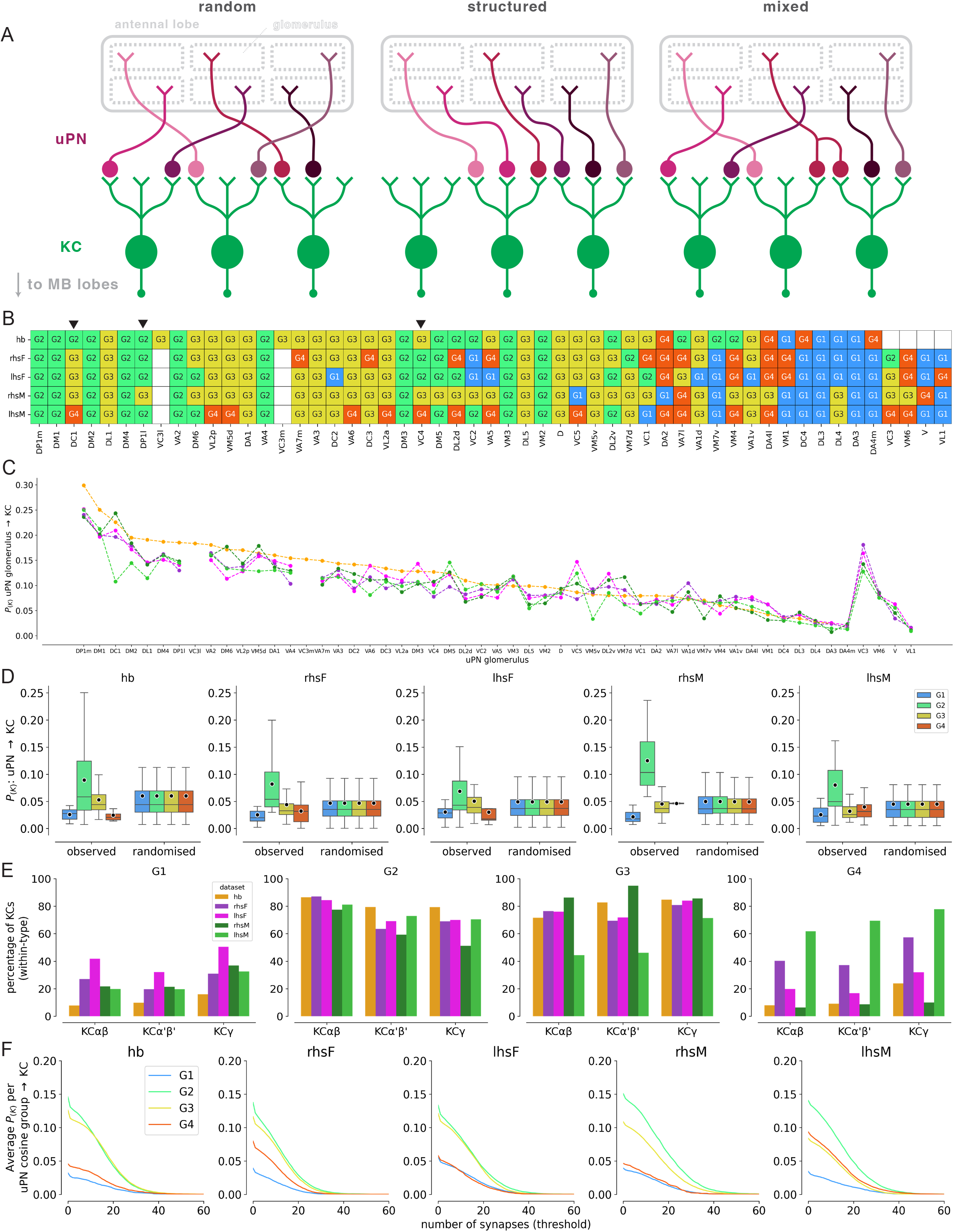
Connectivity clustering reveals biased uPN→KC connectivity in the Calyx: **(A)** Illustration of random, structured and mixed connectivity motifs from uPN glomeruli in the antennal lobe to KCs in the Calyx. **(B)** Canonical cosine cluster assignment for each uPN glomerulus across hemispheres, aligned to (C), with black arrows indicating uPN glomeruli that vary in their canonical cluster assignment across hemispheres. **(C)** Distribution of *P_(K)_* for each uPN glomerulus in each hemisphere. **(D)** Permutation test of canonical clusters. Consensus cosine clustering of uPN glomerulus→KC connectivity matrices identified four canonical cosine clusters across the five hemispheres, each with a distinct connection probability (*P_(K)_*) onto KCs. The G2 cluster consistently shows a higher *P_(K)_* than the other clusters across all hemispheres. To test whether this was greater than expected by chance, connectivity matrices were shuffled (×10,000) and *P_(K)_*recalculated for the canonical cosine clusters to generate a randomised null distribution. G2’s observed *P_(K)_* consistently exceeded this null distribution, indicating that its elevated connection probability is unlikely to arise by chance. Boxplots represent the median values (horizontal lines) and interquartile ranges. The black dots represent the mean. **(E)** Percentage of each KC type targeted by the canonical cosine cluster across hemispheres. **(F)** Threshold analysis of the *P_(K)_* distributions for each canonical cosine cluster.

These results indicate a single canonical cluster (G2) conserved across hemispheres, accompanied by three more flexible clusters (G1, G3 and G4) whose membership varied more frequently. Jaccard comparisons confirmed that G2 best matched one original cluster in every hemisphere, whereas the remaining groups were more variable (Supplementary Figure 3.2A). However, there were differences in the glomerular membership of the G2 canonical cluster between hemispheres (Figure 3B, black arrows). Thus, the G2 canonical cluster contains both a conserved set of glomeruli (Figure 3B, green columns) and a subset of glomeruli that sit at the boundaries between canonical clusters, with their membership shifting between hemispheres rather than remaining fixed (Figure 3B, black arrows).

We calculated *P_(K)_*, the proportion of KCs targeted by a given glomerulus, finding a consistent distribution across hemispheres and datasets (Figure 3C). G2 glomeruli had higher probabilities of connecting to a KC (occurring on the left side of Figure 3C), while G1 glomeruli had lower connection probabilities (occurring on the right side of Figure 3C). Therefore, we next tested whether these canonical clusters were more (or less) likely than chance to connect to a KC (Figure 3D). To do this, we compared the observed values of *P_(K)_* with values from clusters with randomly permuted membership (Figure 3D; Supplementary Figure 3.2D; see Methods). The G1, G3 and G4 canonical clusters had connectivity that was indistinguishable from random (random permutation test, *p* > 0.05, all BH-FDR-adjusted; Figure 3D; Supplementary Figure 3.2D). However, the G2 glomeruli within the G2 canonical cluster consistently connected onto a higher proportion of KCs than expected by chance across hemispheres (random permutation test, *p* < 0.01, all BH-FDR-adjusted; Figure 3D). Moreover, glomeruli from G1 and G4 canonical clusters connected to fewer KCs than G2 and G3 (Figure 3E), an effect that was robust when weak synapses were excluded (Figure 3F).

To investigate whether the structured connectivity of the G2 canonical cluster conveyed information about odour scenes, we aligned glomerular membership to odour scenes and canonical clusters (Supplementary Figure 3.2E–F). The G2 canonical cluster was dominated by uPN glomeruli encoding “yeasty”, “fruity” and fermentation-related odorants, which are all associated with the detection of food (Supplementary Figure 3.2F). We also noted that the innervation density (cable length per μm^3^ of Calyx volume) of uPN axons strongly correlated with *P_(K)_* (Supplementary Figure 3.3B). However, neither *P_(K)_* nor uPN axonal innervation density in the Calyx correlated with mean normalised connection weight onto KCs (Supplementary Figure 3.3B–D). Thus, food-odour-encoding uPNs devoted greater axonal cable length within the Calyx, enabling more synaptic connections with different KC dendrites, but this increase in cable length did not increase the weight of those connections.

### Feedforward KC→MBON connectivity

If there is structured connectivity in KC input, is there also structured connectivity from KCs to MBONs? KC axons define specific horizontal and vertical lobes and this organisation is respected by the dendrites of most MBON types which have therefore long been known to receive input preponderantly from subsets of the five KC types already mentioned; indeed the unified nomenclature for MBONs is based on the lobes occupied by their dendrites (Aso *et al*., 2014a). However, as noted above, within each lobe there are further KC subtypes with more subtle morphological distinctions (Li *et al*., 2020). We therefore analysed KC-subtype→MBON connections, normalised each by the MBON’s total KC input and averaged the resulting values across the five hemispheres (Figure 4A–D). MBONs allocated KC input differently across cell types (Figure 4A). For KCαβ-innervated MBONs, KCαβ-m usually contributed 50–65% and KCαβ-c 20–35%, whereas KCαβ-p exceeded 10% for only MBON06, MBON19 and MBON14. MBONs receiving KCα′β′ input were typically targeted by all three KCα′β′ subtypes, but with input percentages divided roughly equally between two of the three subtypes (Figure 4A). KCγ-m showed a distinct pattern, dominating most γ MBONs, except MBON27, MBON33 and MBON35, which also received substantial KCγ-d input.

**Figure 4:**
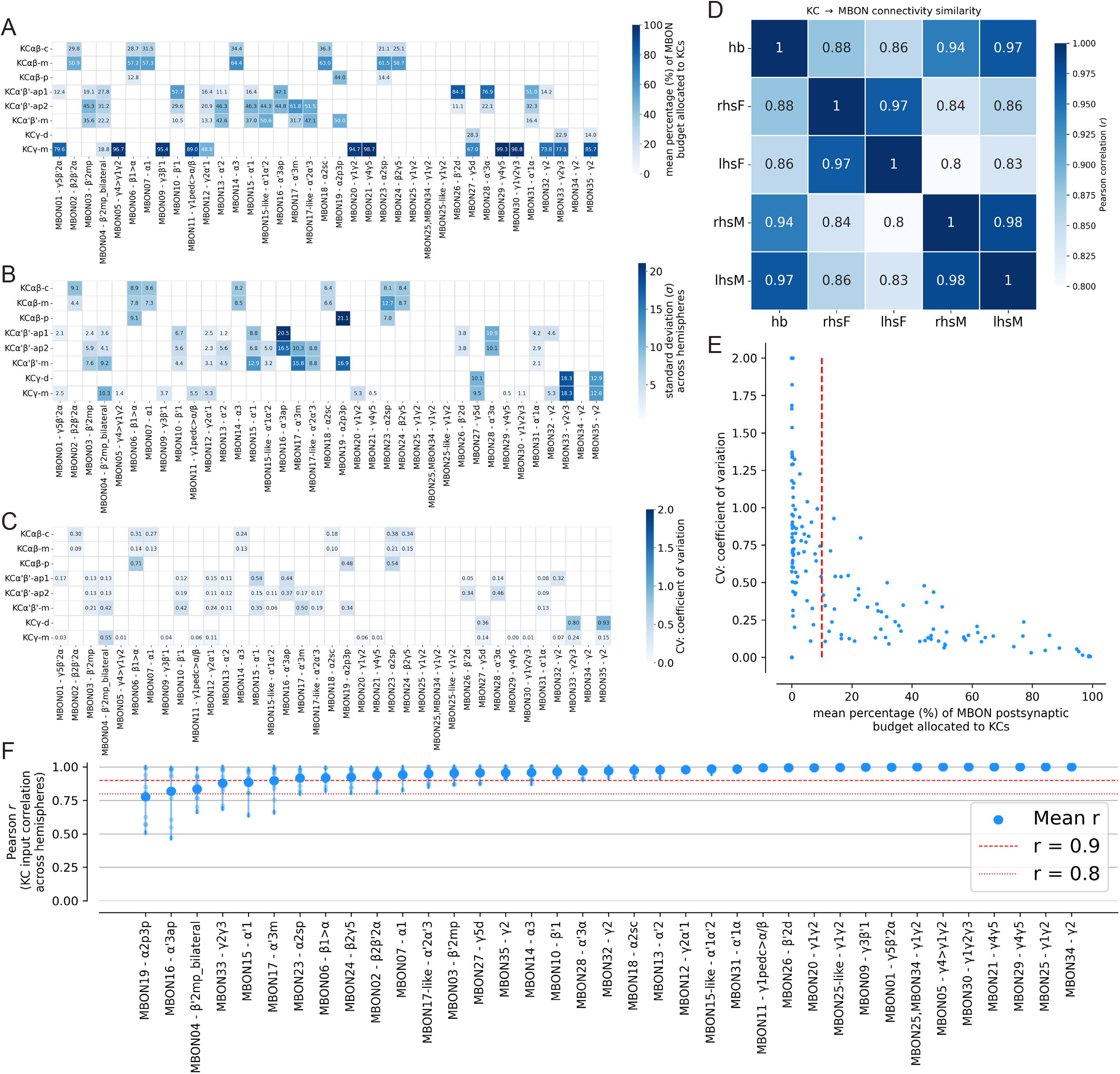
KC→MBON connectivity is largely stereotyped across hemispheres, with variability amongst weaker connections: **(A)** Mean connectivity matrix between KC subtypes and MBON types, created by averaging the connectivity across five hemispheres. A mask was applied to exclude connections <10% of the MBON’s input budget. **(B)** Standard deviation (σ) matrix between KC subtypes and MBON types. Using the mean connectivity matrix in (A), a mask was applied to exclude connections contributing <10% of inputs. **(C)** Matrix of the coefficient of variation (*CV*) for each KC-subtype→MBON-type connection across the five reconstructed hemispheres. *CV* was calculated as the standard deviation divided by the mean proportional input, with higher values indicating greater variability relative to the mean connection strength. **(D)** Pairwise Pearson correlations in KC→MBON connectivity across hemispheres. **(E)** Relationship between mean proportional input and *CV* for individual KC-subtype→MBON-type connections. Each point represents one connection, with the mean percentage contribution to total KC input plotted on the x-axis and *CV* plotted on the y-axis. The red dashed line at 10% indicates the threshold used to distinguish connections with a mean proportional input greater than 10%. **(F)** Per-MBON connectivity correlations in KC inputs across hemispheres. Two red dashed lines indicate Pearson’s *r* values of 0.9 and 0.8, with the mean Pearson correlation value indicated by large blue circles. Paler blue dots within each distribution indicate values from pairwise comparisons for each MBON type.

Most proportional KC-subtype→MBON connections had standard deviations <10% of the MBON’s input, indicating strong conservation (Figure 4B). The main exceptions were KCαβ-p→MBON19 (*σ* = 21.1%) and KCα′β′-ap1→MBON16 (*σ* = 20.5%; Figure 4B). These were both driven by a divergent FlyWire value rather than gradual variation across all hemispheres (Supplementary Figure 4.1). The coefficient of variation (*CV*) declined as mean input weight increased: weak connections varied more proportionally, whereas strong connections were more stable (Figure 4E). Although KCαβ-p→MBON19 and KCα′β′-ap1→MBON16 had high absolute variability, their large means yielded only moderate *CV*s (0.48 and 0.44; Figure 4C). In contrast, KCγ-d→MBON33 and KCγ-d→MBON35 showed the reverse pattern: low means and modest standard deviations produced the largest *CV*s (0.8 and 0.93; Figure 4C). This variation was again FlyWire-specific: in FlyWire, increased KCγ-d input was accompanied by reduced KCγ-m input (Supplementary Figure 4.1).

Overall, KC→MBON matrices were strongly conserved (Figure 4D). KC→MBON connectivity correlations were highest between hemispheres of the same animal (Pearson’s *r* ≥ 0.97; Figure 4D) and lower, though still strong, between animals (Pearson’s *r* = 0.80–0.86; Figure 4D). The majority of MBONs (84%, 31/37) had stereotyped KC-type inputs across all five MBs (Pearson’s *r* ≥ 0.9; Figure 4F), but six MBON types (MBON19, MBON16, MBON04, MBON33, MBON15 and MBON17) had less highly stereotyped KC-subtype inputs (Pearson’s *r* < 0.9; Figure 4F), which may result from biological variation or sample processing.

We then asked whether MBON types comprising two neurons per hemisphere (“sister MBONs”; Figures 1G and 5C) shared similar KC inputs across datasets. Correlating raw KC input weights revealed two strategies: MBON07, MBON09, MBON12 and MBON14 sampled similar KCs with correlated weights (Pearson’s *r* = 0.65–0.88; Figure 5D–E), whereas MBON15, MBON15-like and MBON19 had uncorrelated KC inputs, indicating that these sister MBONs sampled independent sets of KC inputs (Pearson’s *r* = −0.12–0.08; Figure 5D–E). Despite these two groupings, all sister MBONs had highly correlated effective input from uPN glomeruli, as measured by effective uPN-glomerulus→MBON connectivity—the matrix product of the normalised uPN glomerulus→KC and KC→MBON connections (Pearson’s *r* = 0.38–0.99; Supplementary Figure 5.1A–B). Thus, non-overlapping KC sets sampled by each sister cell still converged on the same uPN glomeruli in the Calyx. These data show that MBON wiring can be reliably described and quantified across datasets, and that sensory input from uPNs is stereotyped across sister MBONs, even when fine-scale connections with individual KCs are distinct.

**Figure 5:**
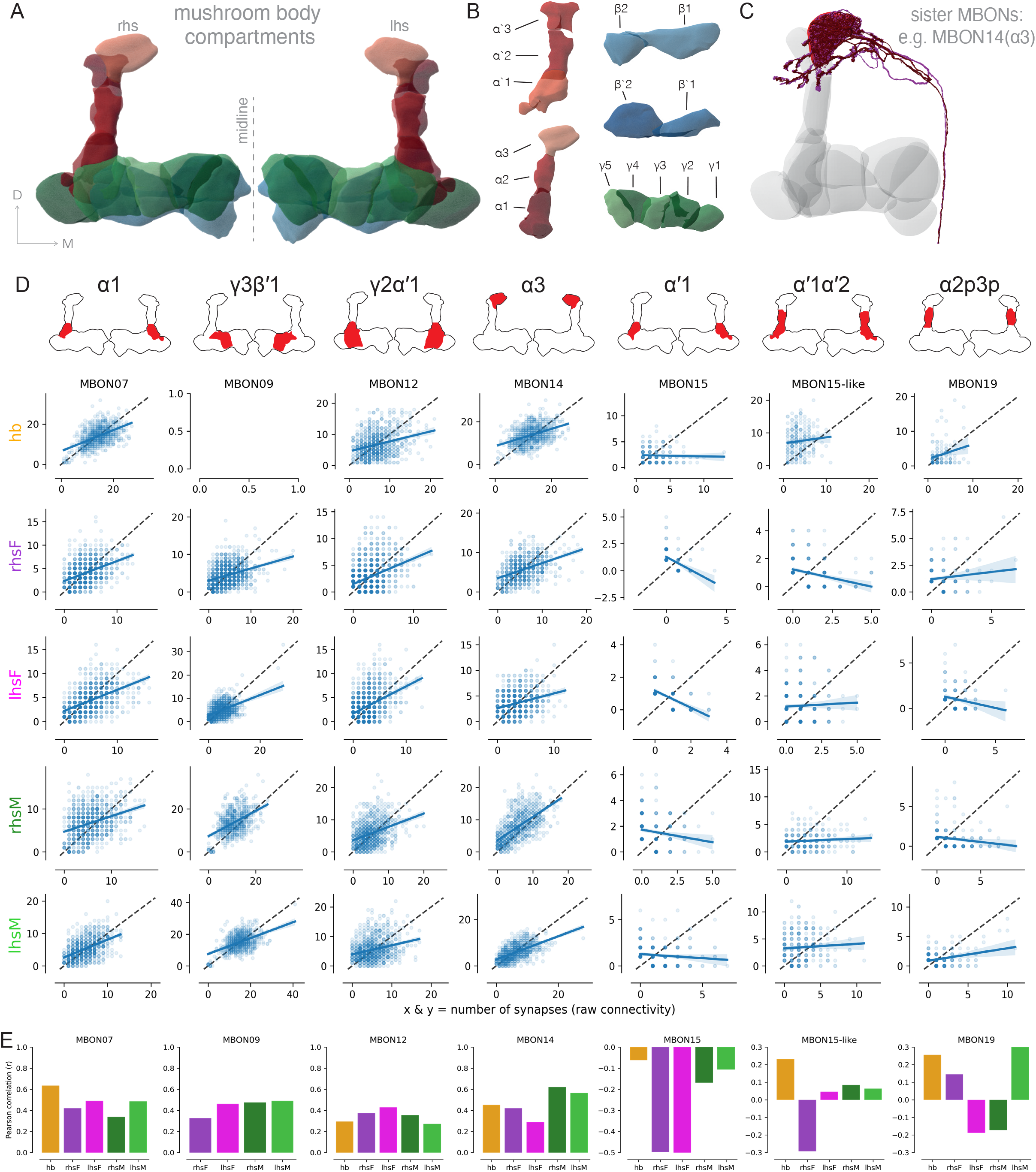
Sister MBON connectivity comparisons reveal different sampling modes of KC input: **(A)** Mushroom Body (MB) compartments. **(B)** Individual MB compartments. **(C)** FlyWire rendering of an example sister MBON, MBON14 (α3), which has two neurons within the type. **(D)** Linear regression analysis of raw synaptic KC connectivity onto sister MBONs for each hemisphere. Black lines are the identity line (x = y) and blue lines are the lines of best fit from linear regression analysis. Blue envelopes represent 95% confidence intervals. **(E)** Pearson correlation coefficients for each sister MBON pair, across hemispheres, presented in (D).

### Effective connectivity mapping reveals an associative olfactory fovea

Is olfactory space uniformly sampled across MBONs or biased towards ecologically important channels? Given the food-odour biases in uPN glomerulus→KC connectivity and compartmentalised KC→MBON wiring (Figures 3–4), we hypothesised that effective sensory input to MBONs would be non-uniform. We therefore extended our effective uPN glomerulus→MBON analysis to the full MBON population. We used dimensionality reduction (UMAP) to visualise the normalised uPN glomerulus→KC (Figure 6A, first column) and KC→MBON (Figure 6A, second column) connectivity matrices, then multiplied these matrices to calculate effective uPN→MBON connectivity via KCs. We then examined the resulting network from both uPN (Figure 6A, third column) and MBON (Figure 6A, fourth column) perspectives, across all hemispheric MBs (Figure 6A, rows).

**Figure 6:**
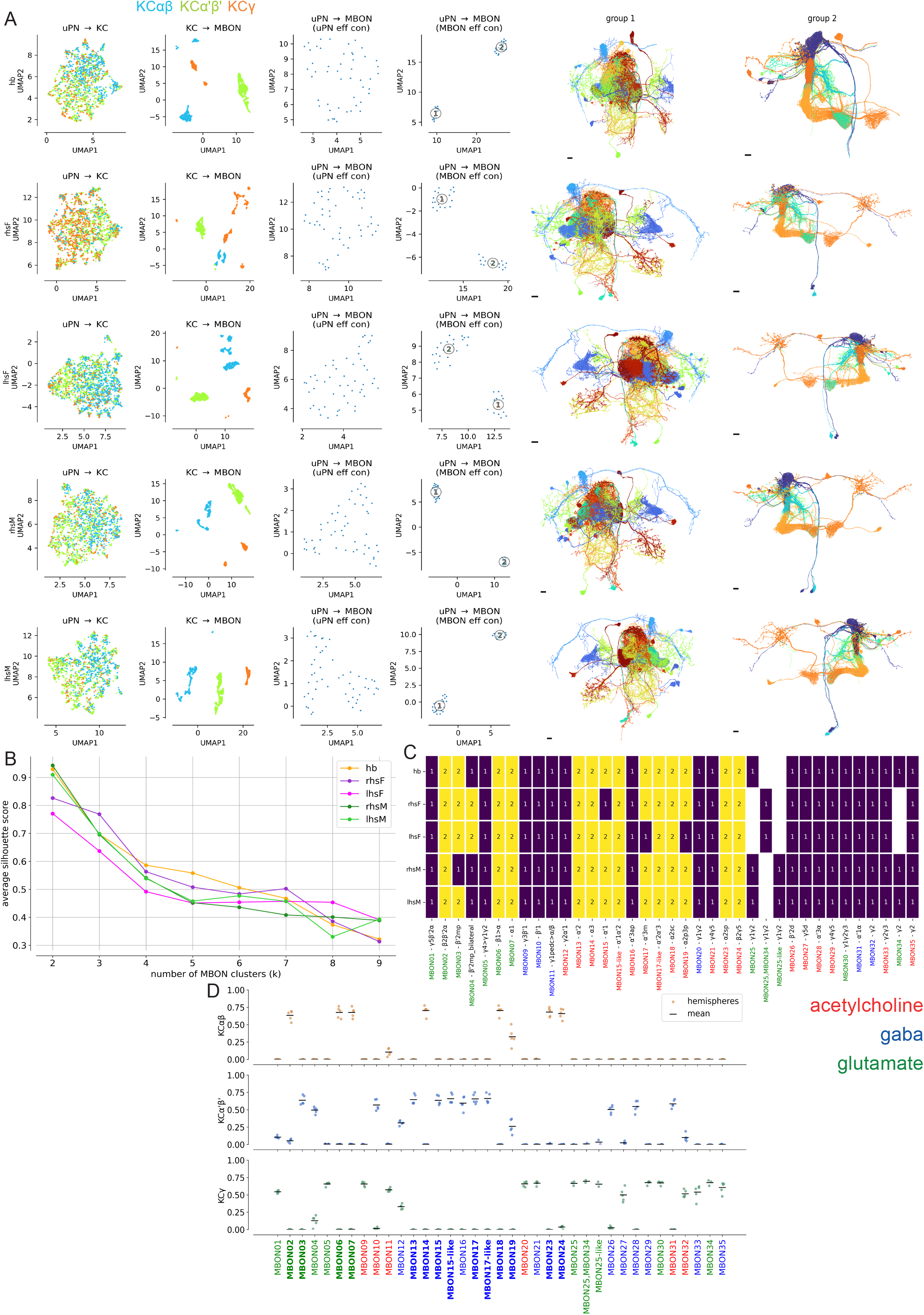
uPN odour channels into the MB reproducibly bifurcate MBONs: **(A)** UMAP embeddings of uPN glomerulus→KC connectivity, KC→MBON and uPN→MBON effective connectivity, with the latter computed from both the uPN and MBON perspectives (i.e. using the effective connectivity matrix and its transpose). The final two columns show MBON morphologies separated by their assigned group and coloured by MBON type. **(B)** Silhouette analysis identifying the optimal number of clusters in the UMAP embeddings of uPN glomerulus→MBON effective connectivity, computed using the transpose (i.e. the MBON-input perspective). **(C)** Allocation matrix showing which group each MBON was assigned to within the UMAP embedding. **(D)** Cross-hemisphere reproducibility of KC-type-specific effective connectivity contributions to MBON types, with MBONs coloured by known and predicted neurotransmitters. For each MBON and each KC type (top: KCαβ; middle: KCα′β′; bottom: KCγ), coloured points show the summed uPN-effective connectivity in each of the five hemispheres analysed, with black lines representing the mean across hemispheres. MBONs in bold indicate MBONs that were allocated to group 2 in the UMAP analysis above.

Clustering analysis of uPN glomerulus→MBON effective connectivity consistently identified two distinct sets of MBONs (Figure 6A). One of these clusters had strong effective connectivity with G2 food-odour uPN channels, notably the DM1 and DP1m glomeruli (Supplementary Figure 6.1). To support the choice of clusters, we compared the similarity of data points within and between clusters using silhouette analysis, which confirmed that two clusters best described the data across all MB hemispheres (Figure 6B). These effective-connectivity data revealed a bifurcation in the pathways linking uPN input to MBONs. Information from food-odour-related glomeruli was routed onto a set of MBONs (group 2 in our clustering; Figure 6C), predominantly by KCαβ and KCα′β′ cell types (Figure 6D; Supplementary Figure 6.2). Group 2 contained predominantly cholinergic, approach-promoting MBONs in the vertical lobe of the MB (Figure 6A—final two columns; Aso *et al*., 2014a; Aso *et al*., 2014b). This conserved non-random structure suggests a specific allocation of neuronal resources for food-odour memory—an ‘associative olfactory fovea’ analogous to that of the visual system.

### Recurrent LHCENT connections target food-odour-associated KCs

Finally, we analysed LHCENT connectivity to understand how MB-driven feedback is aligned with KC inputs (Figure 7). These cell types have dendrites in the superior protocerebral neuropils (SLP, superior lateral protocerebrum; SIP, superior intermediate protocerebrum; SMP, superior medial protocerebrum). It was previously noted that these cell types send feedback to the Lateral Horn (Bates *et al*., 2020), however, we noticed that a subset sends axons to the Calyx (types 1–5 and 8; Figure 7A). Most LHCENT types were predicted to be inhibitory: most were GABAergic; LHCENT4, LHCENT12 and LHCENT14 were glutamatergic (an inhibitory neurotransmitter in the central fly brain; Liu and Wilson, 2013); and LHCENT11 was predicted to be cholinergic (Supplementary Figure 7.1A).

**Figure 7:**
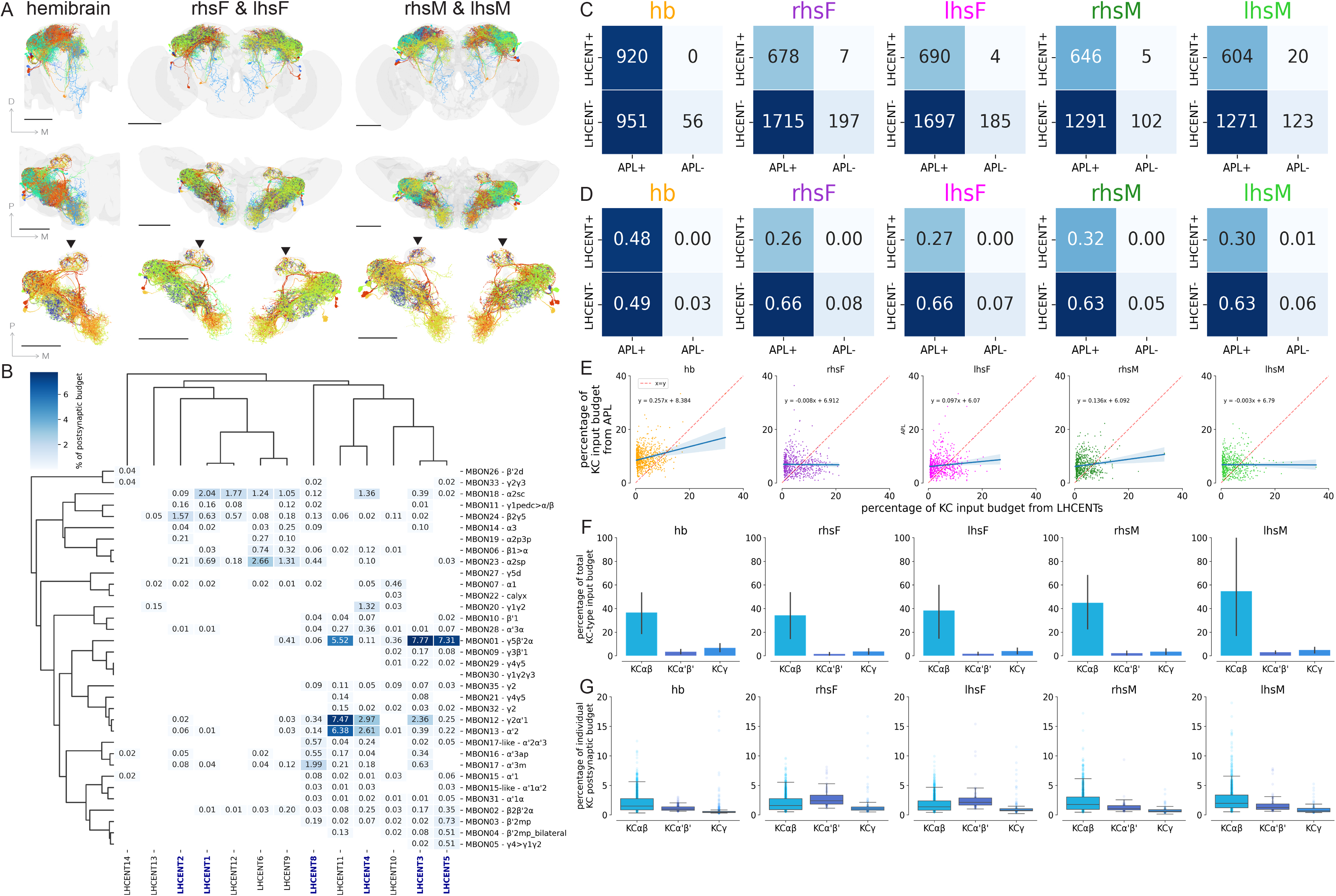
LHCENTs form a recurrent feedback loop from MBONs to KCs: **(A)** Frontal and dorsal views of recurrent LHCENT morphologies in hemibrain, FlyWire and mCNS datasets. The scale bar is 100 μm. Arrowheads indicate axonal projections in the Calyx. **(B)** Averaged MBON→LHCENT connectivity matrix, normalised by the total number of postsynapses each LHCENT has in the SLP/SIP/SMP neuropils. This connectivity matrix was clustered using cosine similarity with WPGMA linkage. LHCENTs that have axonal projections in the Calyx are highlighted in bold. **(C)** The number of KCs receiving input from LHCENTs, APL, both or neither in the Calyx, across hemispheres. **(D)** The proportion of KCs receiving input from LHCENTs, APL, both or neither in the Calyx across hemispheres. **(E)** Connectivity correlations for KCs receiving both LHCENT and APL input in the Calyx (i.e. LHCENT+ and APL+ KCs), normalised by the total number of inputs each KC receives in the Calyx and expressed as percentages. Red dashed lines indicate the identity line (x = y) and blue lines indicate regression lines, with blue envelopes indicating 95% confidence intervals for the regression lines. **(F)** The percentage of total KC-type inputs from all LHCENTs. **(G)** The percentage of input per individual KC from an LHCENT. Each point represents one LHCENT→KC connection; boxes show the median and interquartile range and whiskers extend to the most extreme values within 1.5× the interquartile range.

Across datasets, there were no hemispheric differences in MBON→LHCENT connections in the SLP/SIP/SMP neuropils, normalised by the total number of postsynapses of each LHCENT type in these regions (Kruskal-Wallis test, *H* = 1.56, *p* = 0.815; Supplementary Figure 7.1C). There were also no hemispheric differences in the number of proofread postsynapses (Supplementary Figure 7.1E–J). Therefore, we averaged MBON→LHCENT connectivity across MB hemispheres to enable robust analysis of its circuit structure (Figure 7B). Averaged across all MB hemispheres, individual MBON→LHCENT connections typically accounted for ≤7% of an LHCENT’s input (Figure 7B). However, MBONs could collectively account for up to 20.9% of an individual LHCENT’s total postsynaptic input (Supplementary Figure 7.1D). Individual LHCENTs integrated input across distinct MB lobes and compartments: for example, LHCENT3 received input from MBON01 (γ5β′2a) and MBON12 (γ2α′1), while LHCENT4 integrated MBON12 (γ2α′1) and MBON13 (α′2) (Figure 7B). Thus, LHCENTs integrated inputs from multiple MBON types and compartments, including types that drive both approach and avoidance, rather than being driven by a single output channel.

The LHCENT types 1–5 and 8 had presynapses in the Calyx (Figure 7A), where their primary downstream targets were KCs (Supplementary Figure 7.2A). We examined the number and proportion of KCs co-targeted by both LHCENTs and the anterior paired lateral (APL) neuron, the primary mediator of global inhibition in the MB, which decorrelates KC odour-evoked activity to produce the MB’s characteristic sparse odour code (Lin *et al*., 2014; Figures 7C–D). Approximately 50% of KCs in hemibrain received dual inhibition from LHCENTs and APL, compared with ∼22–23% in FlyWire and ∼27–30% in mCNS (Figure 7D). As such, the fraction of the KC population subjected to convergent localised and global inhibition varied by more than twofold across these hemispheres. At co-innervated KCs, APL→KC and LHCENT→KC connection weights were only weakly but positively correlated, and LHCENT input was consistently weaker than APL input (Figure 7E; Supplementary Figure 7.2B). This pattern suggests that LHCENTs and APL make parallel, distinct inhibitory contributions to KCs.

Is LHCENT inhibition targeted towards specific KC populations? To explore this, we first examined which KC cell types are primarily innervated by LHCENTs in the Calyx (Figure 7F). Within each MB hemisphere, we summed the synaptic input supplied by LHCENTs to each KC cell type (KCαβ, KCα′β′ and KCγ) and normalised this value by the total input received by that KC cell type (Figure 7F). Consistently across datasets, KCαβ received the largest proportional LHCENT input (Figure 7F). Then, to examine how this input to particular KC cell types was distributed across individual KC partners (Figure 7G), we normalised every LHCENT→KC connection by the total synaptic input received by the postsynaptic KC in the Calyx. Median fractional LHCENT input was highest for KCαβ in hemibrain (1.49%) and both mCNS hemispheres (1.97% and 1.75%), whereas KCα′β′ had the highest medians in the two FlyWire hemispheres (2.13% and 2.38%; Figure 7G). KCγ consistently had the lowest median input, ranging from 0.45% to 0.99% across datasets. Therefore, LHCENTs predominantly, but not exclusively, targeted KCαβ neurons in the Calyx. Among LHCENT→KC connections, fractional input weights were broadly comparable between KCαβ and KCα′β′, whereas KCγ consistently received lower median fractional input.

The observation that LHCENTs predominantly targeted KCαβ neurons aligns with our earlier finding that G2 food-odour-encoding uPN glomeruli consistently targeted similar numbers of KCαβ across hemispheres (Figure 3E). To explore the alignment of LHCENT feedback with KCs driven by food-odour-related uPN input, we calculated the similarity between the KC targets of uPN glomeruli and LHCENT types (Supplementary Figure 7.2C). LHCENTs preferentially targeted the same individual KCs that received inputs from the G2 food-odour uPN glomeruli (e.g. DM1, DM2 and DP1m; Supplementary Figure 7.2C–D). The MB therefore combines dense food-odour sampling with an MBON-driven, recurrent GABAergic LHCENT feedback loop. By preferentially targeting KCs receiving food-odour-enriched input, this feedback pathway may gate activity within the MB’s ‘associative olfactory fovea’, where sensory information is preferentially routed towards memory formation.

## Discussion

Across five MB hemispheres from three adult *Drosophila melanogaster* connectomes, we found stereotyped circuit organisation despite variation in cell number and finer-scale connectivity. uPNs, most MBONs and LHCENTs were numerically conserved, whereas KC abundance varied, most notably through KCγ expansion in FlyWire, which is proposed to originate from starvation of that animal in the larval stage (Schlegel *et al*., 2024). Despite this variation, uPN→KC connectivity contained a reproducible non-random bias towards higher-order organisation, and effective connectivity routed these enriched sensory channels preferentially towards a subset of MBONs associated with behavioural approach. MB output, in turn, returns to the Calyx through a recurrent LHCENT pathway that preferentially targets KCs receiving these same food-associated inputs. Importantly, we did not find evidence for sexual dimorphism in the cell types, numerical stereotypy, connectivity budgets or circuit organisation at any stage of the MB network, strongly indicating that the MB is isomorphic. Together, these observations suggest that the MB is neither uniformly random nor rigidly hard-wired. Rather, stable circuit-level organisation is maintained despite substantial variation in cell number and fine-scale connectivity.

### A stereotyped scaffold accommodates variation in KC number and connectivity

KCαβ neurons showed the strongest KC stereotypy: their abundance, total input and uPN-derived input remained similar across hemispheres, and G2 glomeruli recruited comparable numbers of KCαβ neurons (Figure 1E–F and 3E). KCαβ formed a particularly stable component of an otherwise variable KC population. By contrast, FlyWire contained 54–81% more KCγ neurons, especially KCγ-m, plus a smaller KCα′β′ increase. Similar KC numbers in hemibrain and mCNS argue against a simple sex effect and suggest biological variation in KC abundance, potentially linked to developmental history (Lin *et al*., 2013; Schlegel *et al*., 2024). Three animals, however, cannot define the population range.

The addition of the mCNS also challenges a proposed KCγ ‘dilution’ model (Schlegel *et al*., 2024). Despite hemibrain-like KCγ numbers, mCNS resembled FlyWire in the number of unique uPN inputs per KCγ (Figure 2C). KC abundance alone therefore cannot explain olfactory convergence. Subtype-specific synaptic budgets or developmental wiring rules may also contribute. Fine-scale variation did not necessarily alter higher-order sensory organisation. Grouping uPN inputs by odour scene revealed similar fractional allocation across KC classes across hemispheres (Figure 2G), suggesting that different synaptic implementations can preserve broad sensory organisation. Technical differences constrain cross-connectome comparisons. Hemibrain and mCNS used FIB-SEM, whereas FlyWire used ssTEM, alongside different processing and synapse-prediction pipelines. Synapse audits revealed systematic recovery differences (Supplementary Figure 2.2), so we emphasise reproducible organisational features and normalised connectivity over absolute synapse counts.

### Structured olfactory input creates an associative fovea

The uPN→KC layer of the MB is often modelled as random (Caron *et al*., 2013; Litwin-Kumar *et al*., 2017), but our data support mixed random-like and structured wiring. Most glomerular groups matched shuffled expectations, whereas G2 contacted more KCs than expected across all hemispheres (Figure 3). Its core membership remained stable despite variation at cluster boundaries. G2 contains many glomeruli tuned to fruity, yeasty and fermentation-related odours, extending evidence for structured food-odour sampling (Zheng *et al*., 2022). We therefore adopt “associative fovea” to describe a broadly distributed input layer in which ecologically salient channels gain disproportionate access to KCs. Greater uPN axonal innervation density correlated with higher KC connection probability, but not greater mean uPN→KC weights. Preferential access may therefore arise through greater anatomical investment in the Calyx, increasing opportunities to contact KC dendrites without strengthening individual connections.

This wiring could contribute to structured KC odour responses. Although KCs have sparse odour responses (Turner *et al*., 2008; Honegger *et al*., 2011; Campbell *et al*., 2013), recent population recordings reveal reproducible tuning to related natural odours, including fruity esters (Yang *et al*., 2023). Preferential uPN convergence offers a plausible anatomical substrate for this reproducible tuning, but physiology must test this link. Biased sampling also occurs in other expansion networks, including *Drosophila* species, the piriform cortex and cerebellar granule-cell circuits (Ellis *et al*., 2024; Pashkovski *et al*., 2020; Chen *et al*., 2022; Nguyen *et al*., 2023). Such biases may preserve combinatorial capacity while improving reliable transmission of behaviourally important signals.

### Feedforward connectivity preserves sensory organisation at MB output

Structured organisation continued downstream of the Calyx. Most MBONs received reproducible KC-subtype input proportions, and stronger KC-subtype→MBON connections varied less across hemispheres than weaker ones (Figure 4). Major feedforward pathways therefore appear more stable than low-weight connectivity.

Sister MBONs used two wiring strategies (Figure 5). MBON07, MBON09, MBON12 and MBON14 sampled overlapping KCs with correlated weights, potentially supporting a reliable redundant readout. This network-level precision resembles, at a different scale, the matched structural strengths of repeated hippocampal synaptic contacts described by Bartol *et al*. (2015). MBON15, MBON15-like and MBON19, however, sampled largely independent KC ensembles, potentially increasing representational capacity. Despite these different KC-sampling strategies, sister MBONs received highly correlated effective uPN input profiles. Fine-scale partner identity can therefore vary while preserving population-level sensory coding, allowing different KC ensembles to implement similar sensory-to-memory transformations. The relationship between anatomical and functional stereotypy is likely to depend strongly on the level at which it is measured. Hige *et al*. (2015) found a cross-animal rank order for MBON18 and MBON19 odour tuning different from that observed anatomically here. This structural-functional dissociation is plausible because activity also depends on synaptic efficacy, modulation, inhibition, intrinsic excitability and convergence from similarly tuned KCs. Both MBONs innervate the α2 compartment, suggesting that this compartment may be a focus of individualisation and plasticity.

Across the full MBON population, effective connectivity reproducibly separated MBONs into two groups (Figure 6). One received stronger input from food-associated G2 channels, mainly through KCαβ and KCα′β′, and included several cholinergic, approach-promoting MBONs (Aso *et al*., 2014b). The food-odour bias established at the Calyx therefore appears not merely to increase representation at the input layer, but to propagate through the MB towards a particular subset of output pathways. This routing strengthens the associative-fovea model: food-associated channels gain preferential access to KCs and then to approach-associated MBONs. Its persistence despite KCγ expansion in FlyWire further suggests that circuit-level output organisation is robust to variation in KC number.

### Recurrent LHCENT feedback may regulate privileged sensory channels

LHCENTs provide a counterpart to this biased feedforward pathway. They integrate MBON input across superior protocerebral regions, and a subset projects back to the Calyx (Figure 7). Neurotransmitter predictions classified most recurrent LHCENTs as GABAergic and LHCENT4 as glutamatergic; glutamate can also inhibit *Drosophila* olfactory circuits (Liu and Wilson, 2013). This feedback differs from the broad inhibition supplied by APL. Co-innervation varied more than two-fold across datasets, and APL→KC and LHCENT→KC strengths correlated only weakly. We found little structural evidence for a fixed compensatory relationship in which individual KCs trade stronger APL input for weaker LHCENT input, or vice versa. Instead, the two pathways appear anatomically consistent with distinct inhibitory operations: APL provides widespread inhibition across the KC population, whereas LHCENTs contact a more restricted subset of KCs.

The identity of those KCs is particularly informative. LHCENTs preferentially innervated KCαβ neurons and co-targeted KCs receiving G2 inputs, including DM1, DM2 and DP1m. The recurrent pathway is therefore directed towards the same sensory channels that are over-represented in the feedforward network. This arrangement suggests a potential mechanism for selectively regulating privileged olfactory inputs after information has passed through the MB output layer. This organisation could provide localised gain control. Preferential food-odour sampling may improve recruitment of associative circuits but risk over-representing these channels. MBON→LHCENT→KC inhibition could constrain that amplification while allowing mnemonic output to regulate subsequent sensory representations. Neurotransmitter validation, recordings and pathway-specific perturbations must test whether LHCENT feedback alters KC sparseness or food-odour learning.

### Conclusions

Overall, MB stereotypy emerges most clearly at the circuit level. Stable uPN, KCαβ, MBON and LHCENT populations coexist with variation in KCγ abundance and fine-scale wiring, while food-associated input biases, output routing and recurrent feedback remain reproducible across connectomes. We propose that the adult MB combines distributed associative coding with structured pathways that prioritise ecologically important information. This architecture may preserve sparse combinatorial coding while reliably representing food-related stimuli and selectively regulating privileged channels. Additional matched connectomes, potentially from other species, together with physiological and behavioural experiments, will be required to determine which components of this architecture reflect hard-wired developmental rules, individual experience or technical variation between datasets.

## Methods

We analysed three adult *Drosophila melanogaster* electron microscopy (EM) connectomes: Hemibrain v1.2.1, a female FIB-SEM hemisphere (Li *et al*., 2020; Scheffer *et al*., 2020); FAFB/FlyWire version 783 with the Princeton Synapse Release, a bilateral female ssTEM brain (Zheng *et al*., 2018; Dorkenwald *et al*., 2024; Schlegel *et al*., 2024; *Yu et al.*, 2025); and mCNS v1.0, a bilateral male FIB-SEM central nervous system (Berg *et al*., 2026). The hemibrain (hb), FlyWire (lhsF and rhsF) and mCNS (lhsM and rhsM) hemispheres were analysed independently, yielding five Mushroom Body (MB) samples. Hemibrain and mCNS connectivity data were retrieved with neuprint-python; FlyWire matrices were built from the Princeton Synapses data frame (https://codex.flywire.ai/api/download?dataset=fafb). All analyses used Python 3.12 in JupyterLab. Neuronal skeletons, meshes and transformations were processed with navis. Analysis code, neuron tables, connectivity matrices, neuroglancer scenes and compartment meshes are available at https://github.com/markuspleijzier/Pleijzier_et_al.

### Cell-type identification and annotations

uPNs, Kenyon Cells (KCs), Mushroom Body Output Neurons (MBONs) and Lateral Horn Centrifugal Neurons (LHCENTs) were matched across connectomes using published annotations (Li *et al*., 2020; Schlegel *et al*., 2024; Berg *et al*., 2026) and manual review where required. uPNs were typed by antennal-lobe glomerulus innervation; dataset-specific annotation differences were reconciled before comparison, including VC3l/VC3m in hemibrain vs. VC3 in FlyWire/mCNS. KC singleton types and difficult MBON25/25-like/34 assignments were checked manually using morphology and γ-lobe projections. APL was taken from existing annotations. Odour-scene labels were assigned to uPN glomeruli from published receptor/ligand annotations (Huoviala *et al*., 2020; Schlegel *et al*., 2021).

### uPN→KC connectivity and distribution analyses

For each hemisphere, individual uPNs were grouped by glomerulus and their Calyx synapses onto each KC were summed to produce a glomerulus × individual KC matrix. For normalised weight analyses, each connection was divided by the total Calyx input of its postsynaptic KC. For each KC, we also calculated K, the number of unique presynaptic uPN glomeruli with at least one synapse (Schlegel et al., 2024). Pairwise differences in mean K were summarised with Cohen’s *d*:

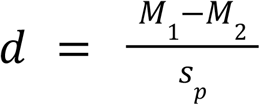

where *s_p_* was calculated as:

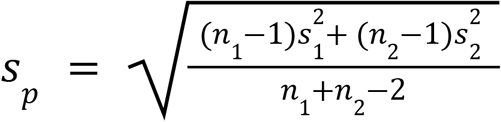

and *n*_1_ and *n*_2_ are the sample sizes of groups 1 and 2, respectively.

Normalised non-zero uPN→KC weights were compared between hemispheres using two-sample Kolmogorov-Smirnov (KS) tests, separately for all KCs and for KCαβ, KCα′β′ and KCγ. *P*-values were corrected with the Benjamini-Hochberg false discovery rate procedure (BH-FDR; α = 0.05); the KS D-statistic quantified effect magnitude. Weight distributions were visualised with Gaussian kernel-density estimation using Scott’s bandwidth rule. To analyse odour scenes, we summed the normalised connection weights from all uPNs assigned to a given odour scene, with KCs receiving no input from that scene assigned a value of zero. For each of nine odour scenes and three KC types, per-KC input distributions were compared across five hemispheres using Kruskal-Wallis (KW) tests (27 comparisons), with BH-FDR correction and epsilon-squared (∊^2^) effect sizes. Significant omnibus tests were followed by pairwise Dunn’s tests with Holm correction, with median differences and Cliff’s delta reported as pairwise effect sizes.

### Synapse auditing

To assess technical variation in synapse recovery, total and proofread pre- and postsynaptic sites were counted within dataset-specific 3D meshes for the Calyx, Peduncle and MB lobes. Proofread connections between major MB cell classes were also tabulated, with a connection defined as an edge between a presynaptic T-bar and a postsynaptic density. The same audit was applied to the SLP/SIP and SMP for LHCENT analyses. These comparisons contextualised differences between FIB-SEM datasets (hemibrain, mCNS) and ssTEM FlyWire.

### uPN→KC clustering, randomisation and structural analyses

To analyse structure in uPN→KC partner selection with minimal assumptions about the nature of the structure, we binarised each glomerulus × KC connectivity matrix (≥1 synapse = 1; no connection = 0) and clustered uPN glomeruli using cosine distance with WPGMA linkage. We derived cross-hemisphere canonical groups by consensus clustering (Monti *et al*., 2003; Strehl and Ghosh, 2002). A co-association matrix recorded the fraction of available hemispheres in which each pair of glomeruli co-clustered; this was converted to distance (1 − co-association) and hierarchically clustered:

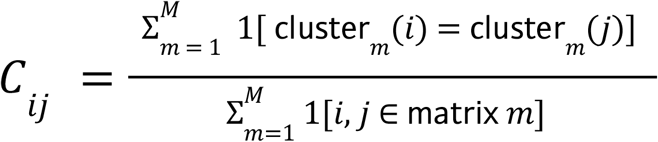

where *M* = 5 is the number of connectivity matrices, and the denominator restricts the comparison to matrices in which both glomeruli *i* and *j* were present, avoiding bias from glomeruli measured in fewer matrices. Cluster labels from each of the five original matrices were mapped onto canonical cluster labels by maximising the Jaccard index 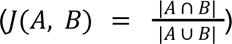 between cluster membership sets.

Hemisphere-specific and canonical groups were matched by Jaccard similarity, allowing many-to-one mappings; low-overlap assignments (*J* < 0.3) were manually reviewed. For each glomerulus, connection probability *P_(K)_*was defined as the proportion of KCs receiving ≥1 synapse. Null distributions were generated from 10,000 row-shuffled matrices per hemisphere, preserving each glomerulus’s connection budget while randomising KC identity. Empirical p-values used a +1 pseudocount and were BH-FDR-corrected:

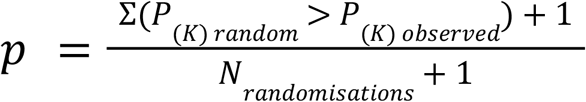

We also calculated the fraction of each KC subtype targeted by each canonical group. To test sensitivity to weak edges, *P_(K)_* was recalculated after incrementally increasing the minimum synapse threshold from 0 to the maximum observed edge weight. uPN innervation density was measured as axonal cable length within the Calyx divided by Calyx volume, and linear regression tested relationships among innervation density, *P_(K)_* and mean normalised uPN→KC weight.

### KC→MBON and effective connectivity

Within each hemisphere, KC→MBON synapses in the MB lobes were grouped by KC subtype and MBON type. For multicellular MBON types, KC inputs were summed across constituent neurons. Each KC-subtype × MBON-type matrix was normalised by each MBON’s total KC input. We calculated the across-hemisphere mean, standard deviation and coefficient of variation 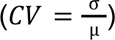 for each connection. Overall matrix similarity and MBON-specific KC-input similarity were quantified with pairwise Pearson correlations across hemispheres. Sister MBONs were compared by correlating raw KC-input vectors for the two constituent cells. Two-step effective uPN→MBON connectivity was calculated for each hemisphere by matrix multiplication of the KC-input-normalised uPN→KC matrix and the MBON-input-normalised KC→MBON matrix. Effective inputs to sister MBONs were compared using Pearson correlation. KC-class contributions were calculated by repeating the multiplication after restricting the KC dimension to KC cell types (KCαβ, KCα′β′ or KCγ).

### Dimensionality reduction

UMAP was used to visualise uPN→KC, KC→MBON and effective uPN→MBON connectivity profiles. Effective connectivity was examined from both uPN and MBON perspectives using the matrix and its transpose. For MBON clustering, UMAP used n_neighbors = 15, n_components = 2 and Euclidean distance. Candidate cluster numbers *k* = 2–10 were compared using mean silhouette scores.

### LHCENT analyses and neurotransmitter predictions

MBON→LHCENT connectivity was restricted to the SLP/SIP/SMP and normalised by each LHCENT’s total input in these neuropils. Hemisphere distributions were compared using empirical cumulative distribution functions (ECDFs) and a KW test; subsequent analyses used the across-hemisphere mean matrix. For Calyx feedback, KCs were classified as receiving LHCENT input, APL input, both or neither, based on a criterion of ≥1 proofread synapse. LHCENT→KC and APL→KC weights at co-targeted KCs were normalised by each KC’s total Calyx input and compared using Pearson correlation. LHCENT input was also summarised by KC subtype, and overlap between LHCENT and uPN targets was quantified with Jaccard similarity. LHCENT fast-neurotransmitter predictions came from FlyWire (Eckstein *et al*., 2024) and mCNS (Berg *et al*., 2026). FlyWire synapse-level predictions were collapsed to neuron-level identities using cleft-score-weighted confidence; mCNS consensus annotations followed Berg *et al*. (2026).

### Software versions

Core package versions used for analysis:

| Package | Version |
| --- | --- |
| pandas | 2.2.3 |
| numpy | 1.24.2 |
| navis | 1.5.0 |
| fafbseg | 3.0.6 |
| neuprint | 0.5 |
| seaserpent | 0.3.1 |
| scipy | 1.10.1 |
| sklearn | 1.3.0 |
| statsmodels | 0.14.1 |
| matplotlib | 3.7.2 |
| seaborn | 0.13.2 |
| tqdm | 4.66.1 |
| umap-learn | 0.5.12 |

---

**Supplementary Figure 1.1:**
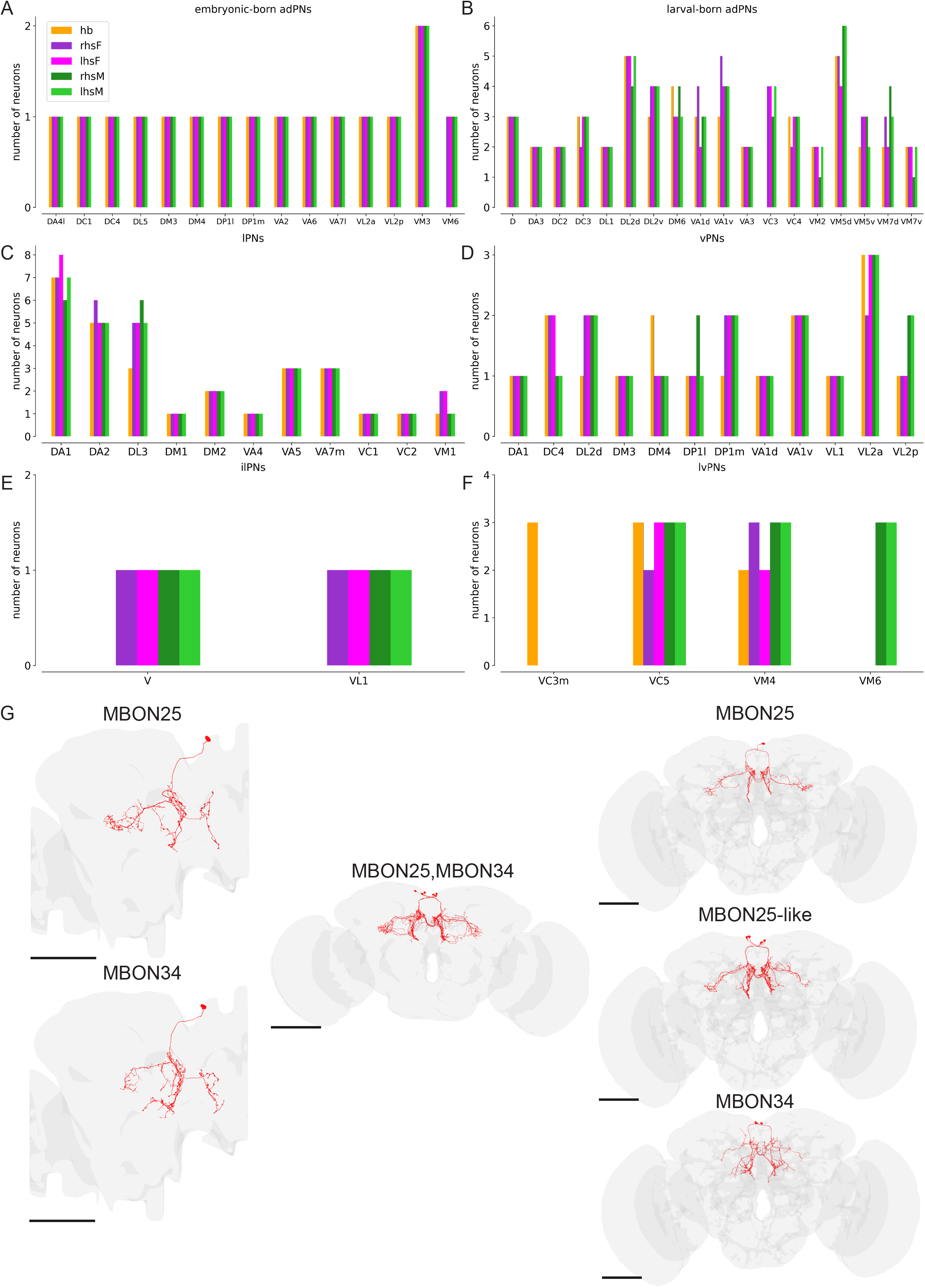
Number of neurons in each **(A)** embryonic-born uPN type, **(B)** larval-born uPN type, **(C)** lPN type, **(D)** vPN type, **(E)** ilPN type, **(F)** lvPN type. **(G)** Morphologies of the difficult-to-type MBONs across hemispheres. The scale bar represents 100 μm.

**Supplementary Figure 2.1:**
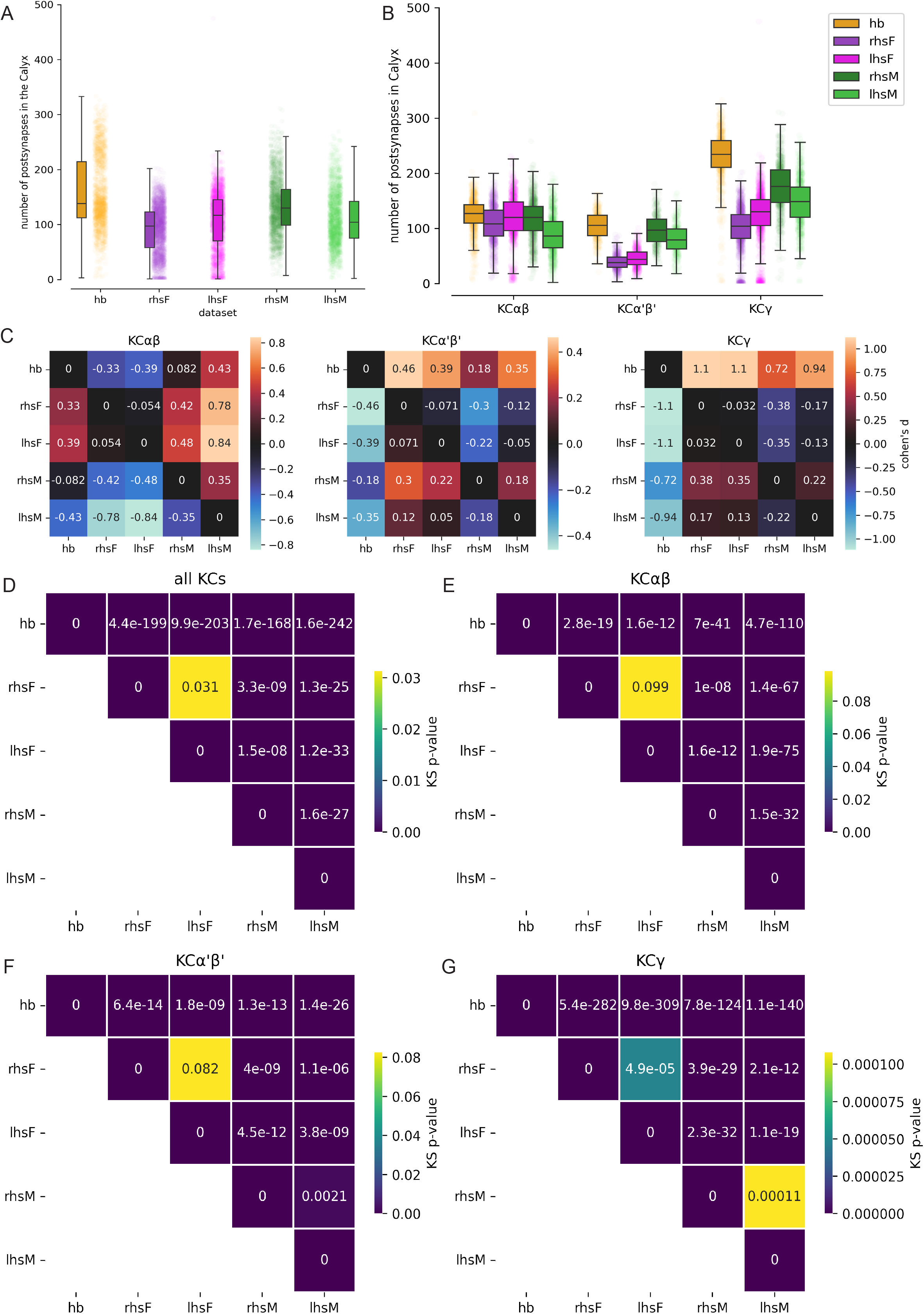
**(A)** Number of postsynapses per individual KC in the Calyx, across hemispheres. **(B)** Distribution of the total number of postsynapses per individual KC neuron, for each KC type in the Calyx, across the different hemispheres. **(C)** Cohen’s *d*-statistic matrix for KCαβ (left), KCα′β′ (middle) and KCγ (right) across the hemispheres. *P*-value matrices for pairwise Benjamini-Hochberg-corrected Kolmogorov-Smirnov tests on the uPN glomerulus→KC connectivity distributions presented in Figure 2E–F, for **(D)** all KCs, **(E)** KCαβ, **(F)** KCα′β′ and **(G)** KCγ.

**Supplementary Figure 2.2:**
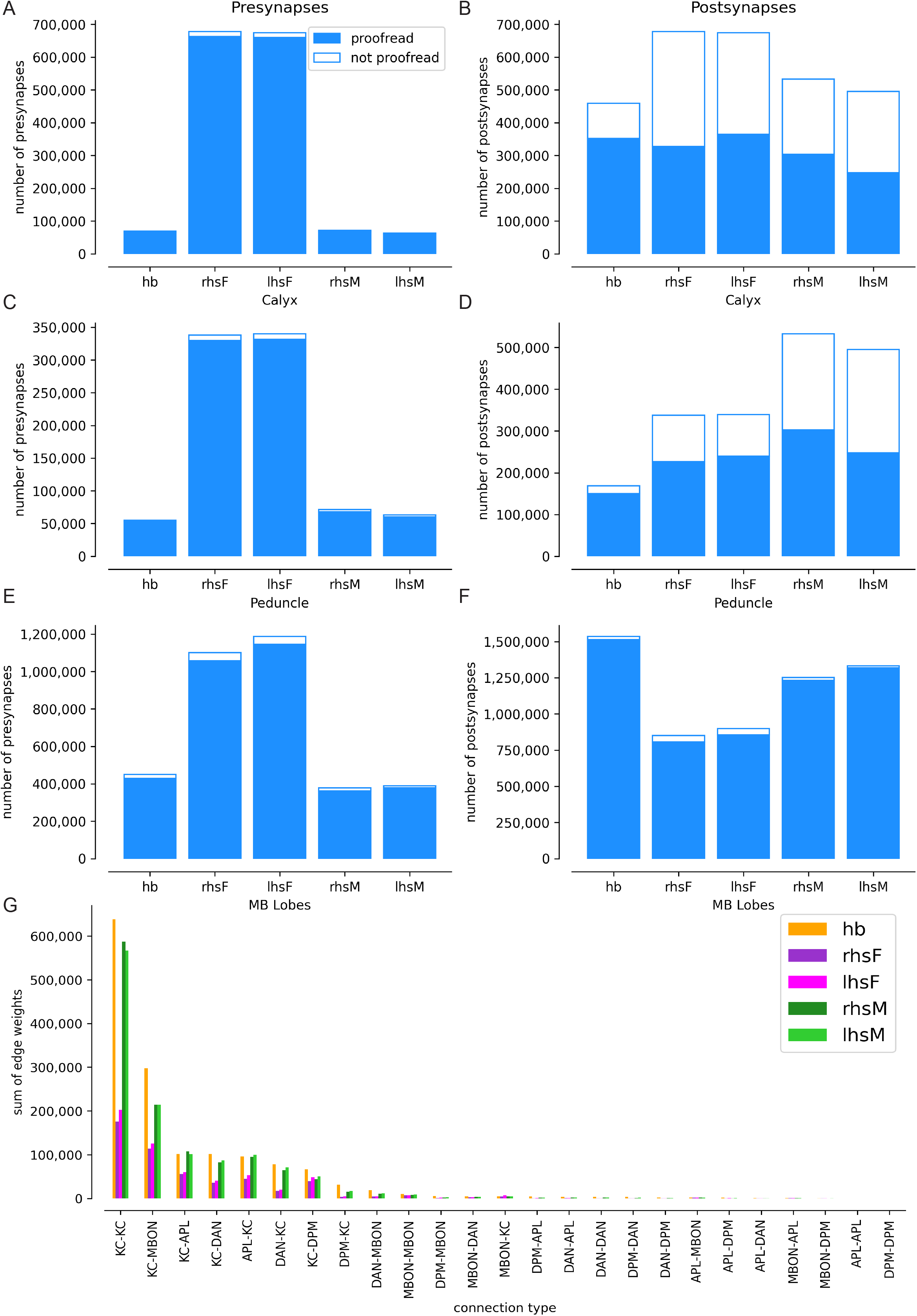
Number of proofread and non-proofread presynapses (left) and postsynapses (right) in the Calyx (**A** and **B**), Peduncle (**C** and **D**) and MB lobes (**E** and **F**). **(G)** Total number of synapses for each connection type in the MB across the five hemispheres.

**Supplementary Figure 3.1:**
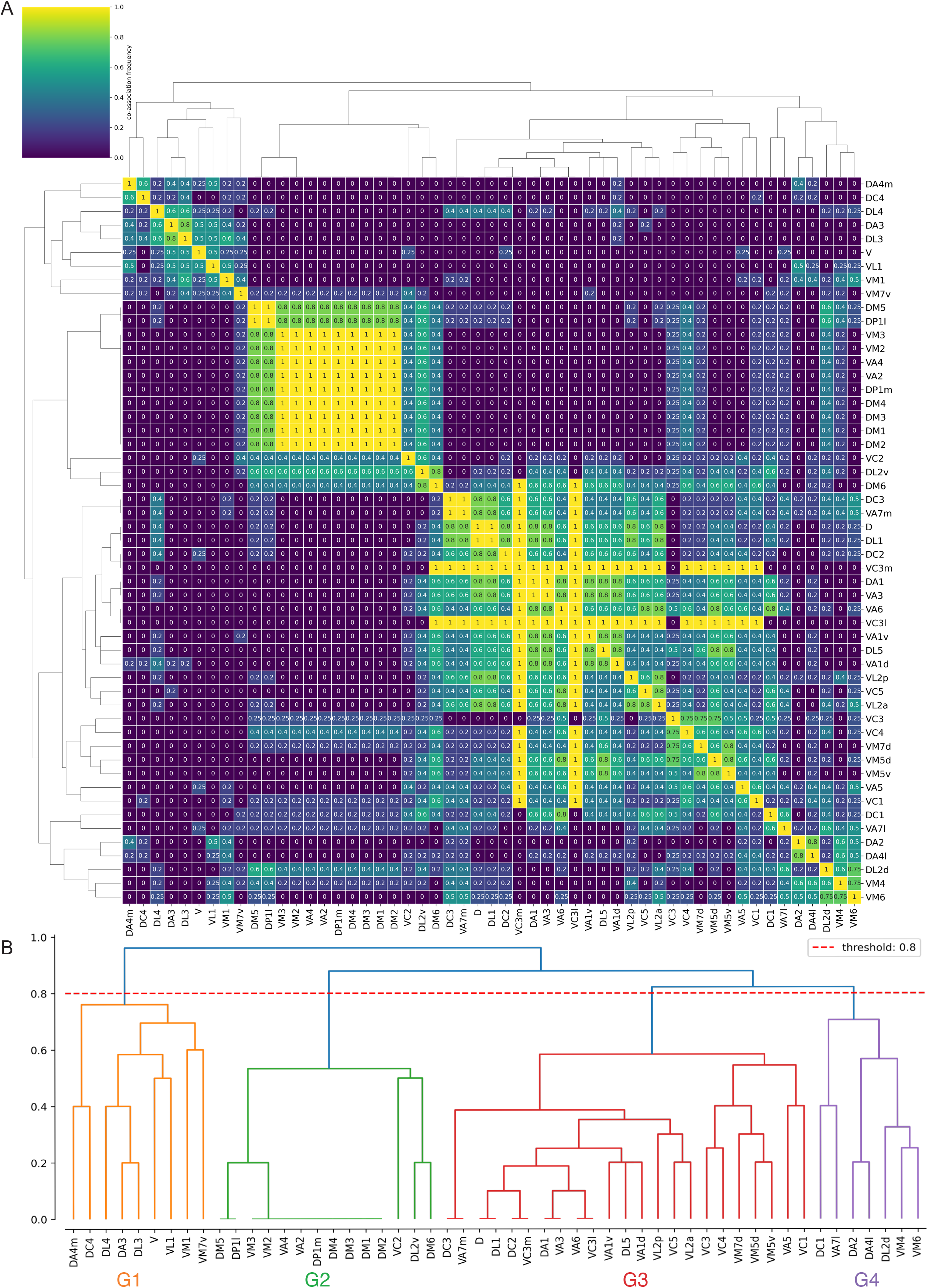
**(A)** Consensus clustering co-association matrix. For each pair of glomeruli present in a given connectivity matrix, we recorded whether the two glomeruli were assigned to the same cluster. Across the five matrices, we constructed a co-association matrix in which each entry represents the proportion of matrices, among those in which both glomeruli were present, that assigned the pair to the same cluster. **(B)** The co-association matrix was converted to a distance matrix (1 − *C*) and subjected to hierarchical clustering using average linkage. The resulting dendrogram was cut at a height chosen by visual inspection to yield canonical clusters.

**Supplementary Figure 3.2:**
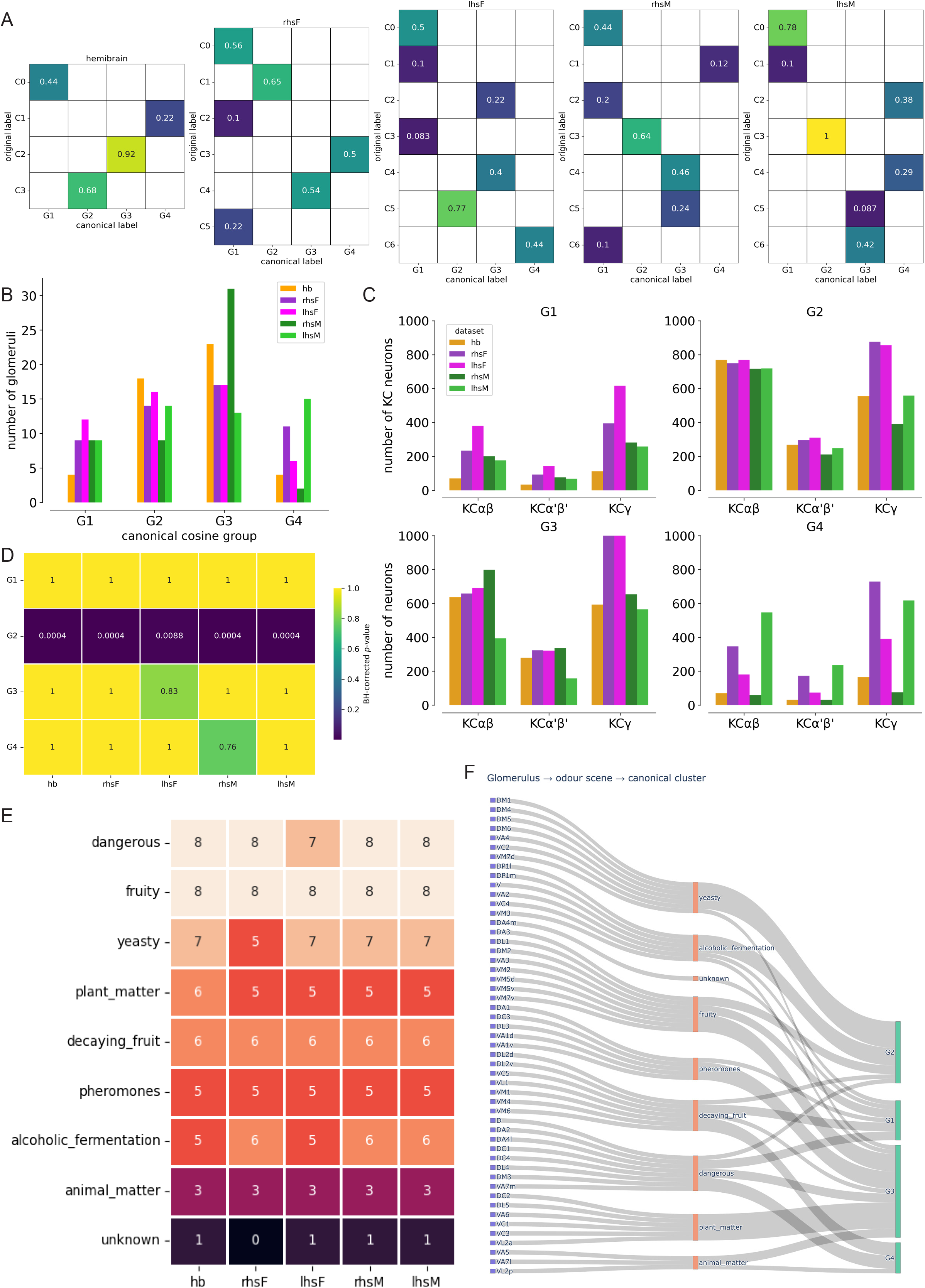
**(A)** Jaccard similarity matrix comparing the original per-hemisphere cosine clusters of uPN glomeruli→individual KC connectivity to the new canonical clusters derived from the consensus clustering approach. **(B)** Number of glomeruli within each of the canonical clusters, across hemispheres. **(C)** Number of individual KCs, by KC type, targeted by each cosine cluster in each hemisphere. **(D)** BH-FDR-corrected *p*-value matrix from the permutation test in Figure 3D, showing that G2’s observed *P_(K)_* exceeded chance expectations (all *p* < 0.01). **(E)** Number of glomeruli within each odour scene. **(F)** Sankey diagram of how each uPN glomerulus maps onto odour scenes, and how each odour scene in turn maps onto the canonical clusters. An interactive version of this diagram is available on this publication’s accompanying GitHub repository (https://github.com/markuspleijzier/Pleijzier_et_al).

**Supplementary Figure 3.3:**
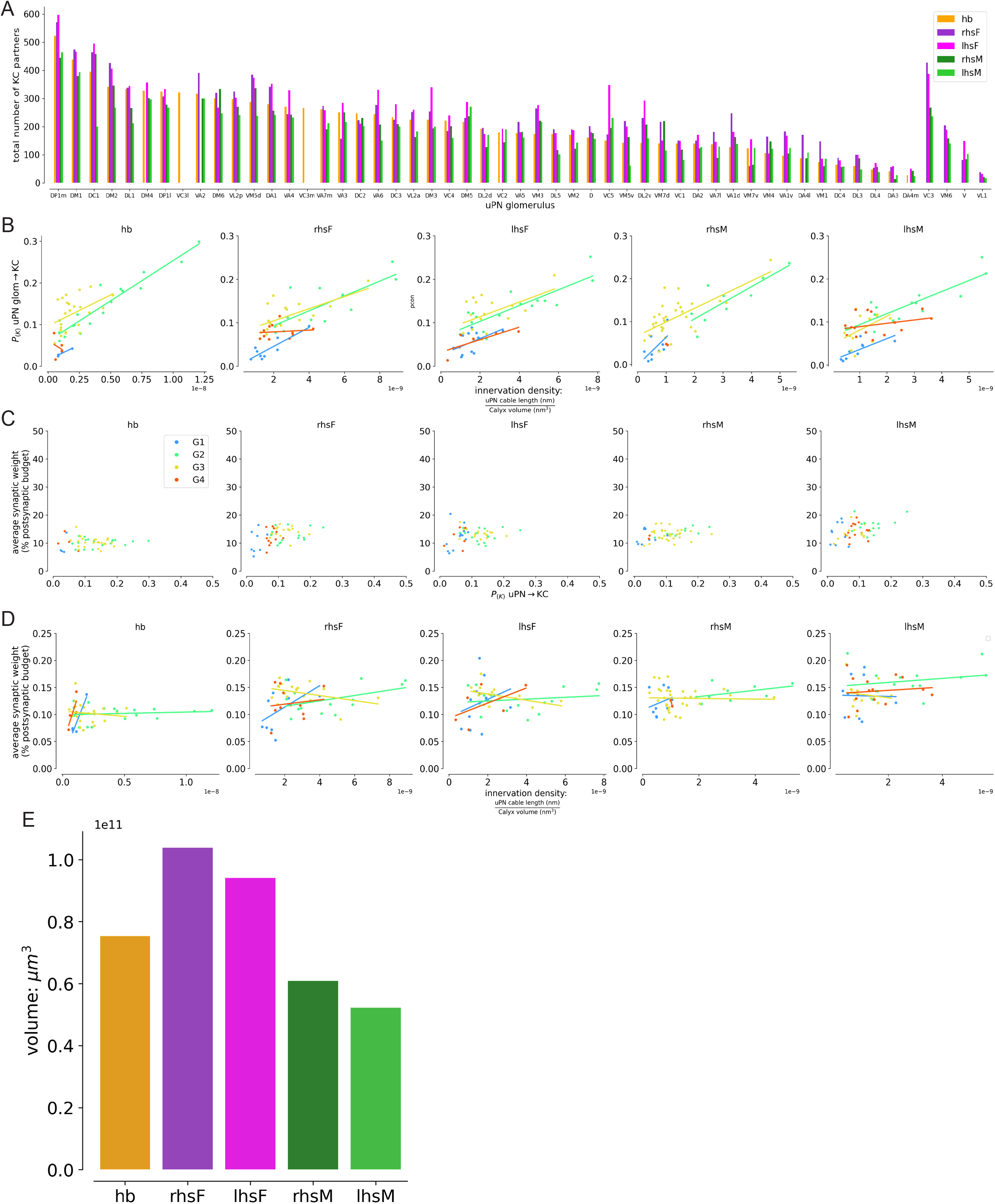
**(A)** Total number of downstream KC partners for each uPN glomerulus across the five hemispheres. **(B)** Correlations between innervation density and *P_(K)_*for each glomerulus, across the five hemispheres. **(C)** Correlations between *P_(K)_* and mean synaptic weight for each glomerulus across the five hemispheres. **(D)** Correlations between innervation density and mean synaptic weight for each uPN glomerulus across the five hemispheres. **(E)** Calyx volume (µm^3^) for each of the five hemispheres, used for calculating innervation density.

**Supplementary Figure 4.1:**
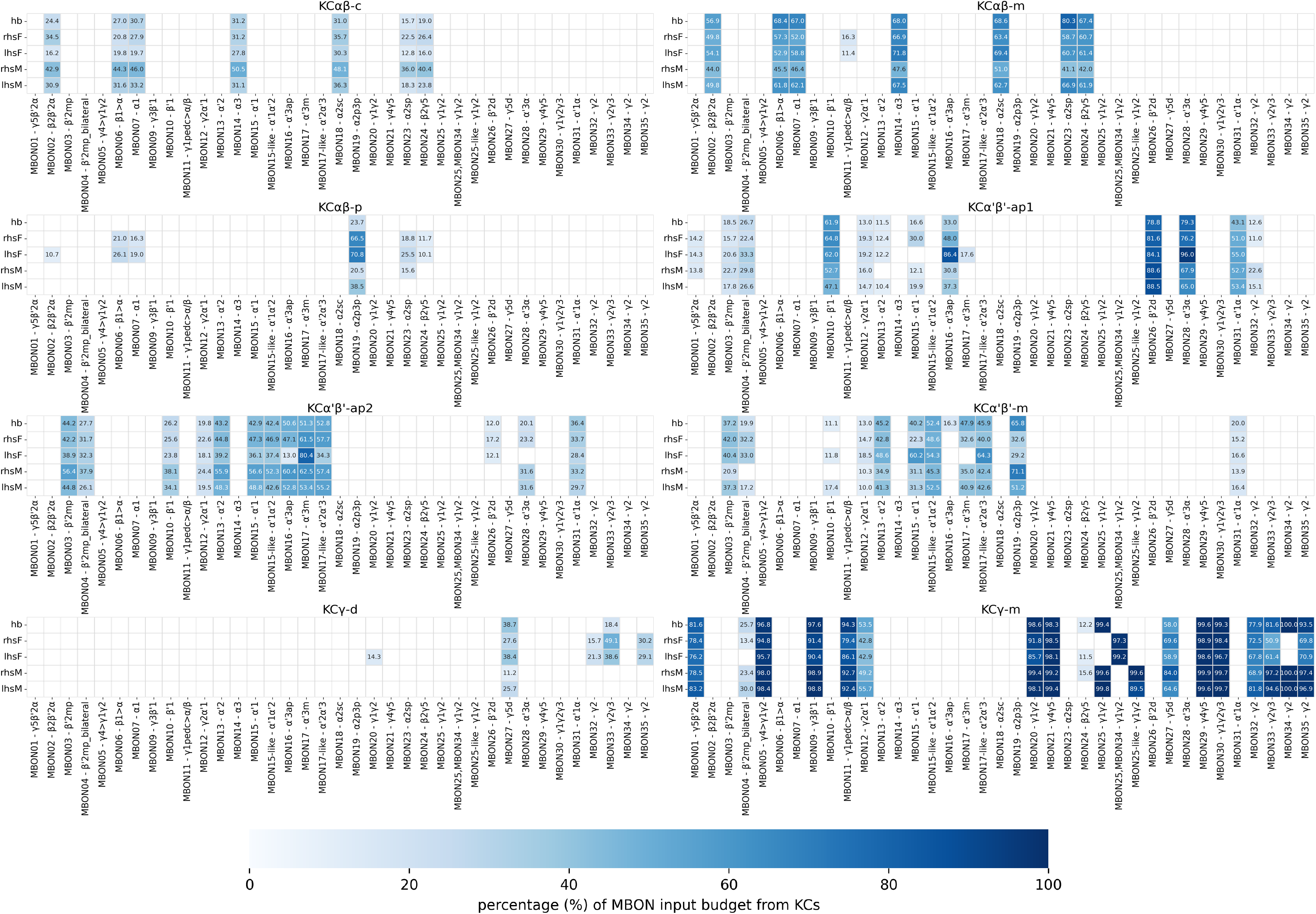
KC-subtype→MBON connectivity matrices across five MBs. Heatmaps show the proportional contribution of eight KC subtypes (KCαβ-c, KCαβ-m, KCαβ-p, KCα′β′-ap1, KCα′β′-ap2, KCα′β′-m, KCγ-d and KCγ-m) to the total KC input received by each MBON type. Each heatmap corresponds to one KC subtype, with rows representing the five reconstructed hemispheres and columns representing the aligned MBON types. For each hemisphere and MBON type, connectivity proportions were calculated by dividing the synaptic input from the indicated KC subtype by the total KC input received by that MBON and displayed as percentages. Higher values indicate that the corresponding KC subtype accounts for a greater proportion of the MBON’s total KC input. MBON types are presented in the same order across all matrices, using the same colour scale for each matrix, allowing for direct comparison between KC subtypes and hemispheres.

**Supplementary Figure 5.1:**
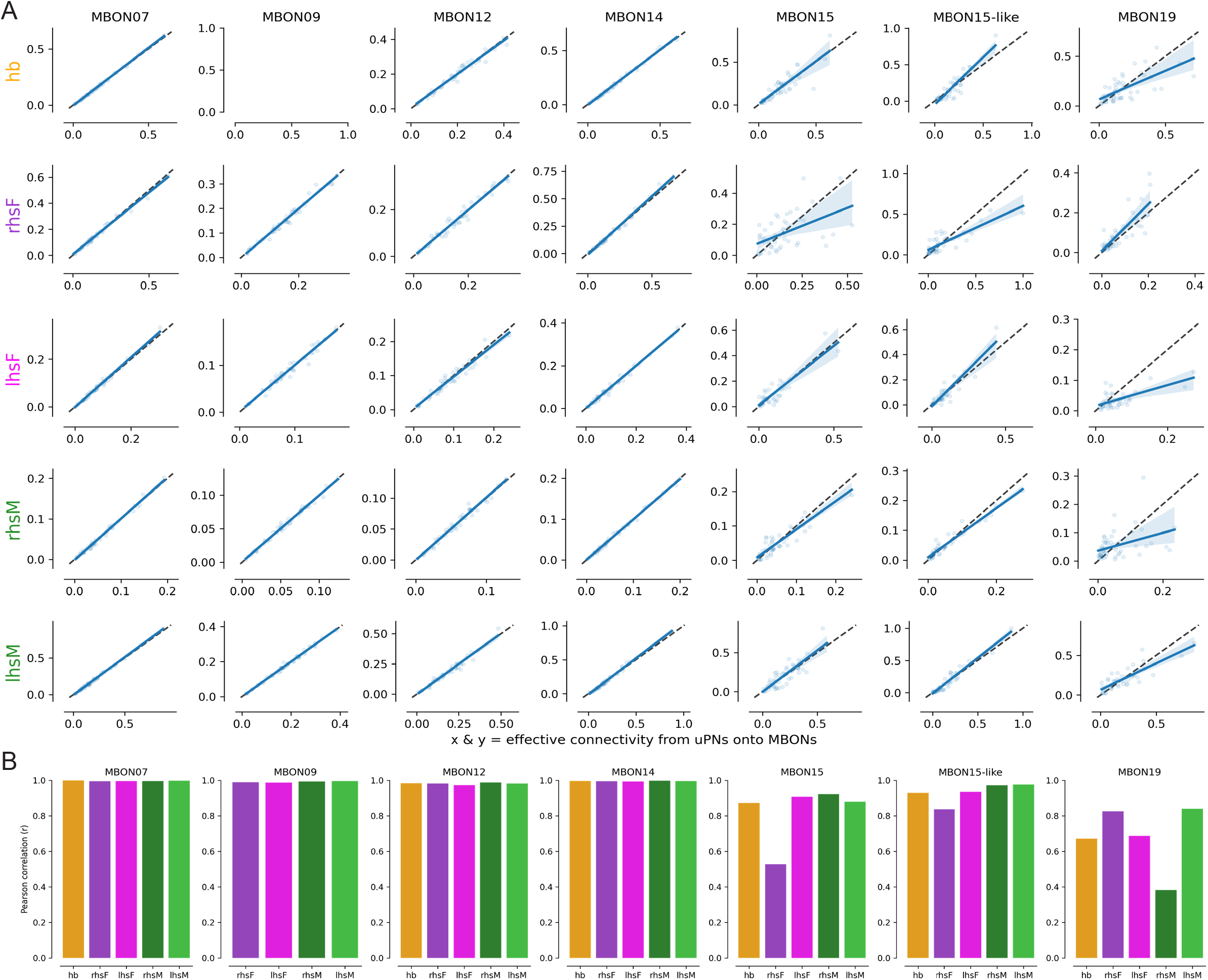
**(A)** Linear regression analysis of the effective uPN glomerulus→MBON connectivity (via KCs) between sister MBONs. Black lines are the identity line (x = y) and blue lines are the lines of best fit. Blue envelopes represent 95% confidence intervals. **(B)** Pearson correlation coefficients for the effective uPN glomerulus→MBON connectivity presented in (A).

**Supplementary Figure 6.1:**
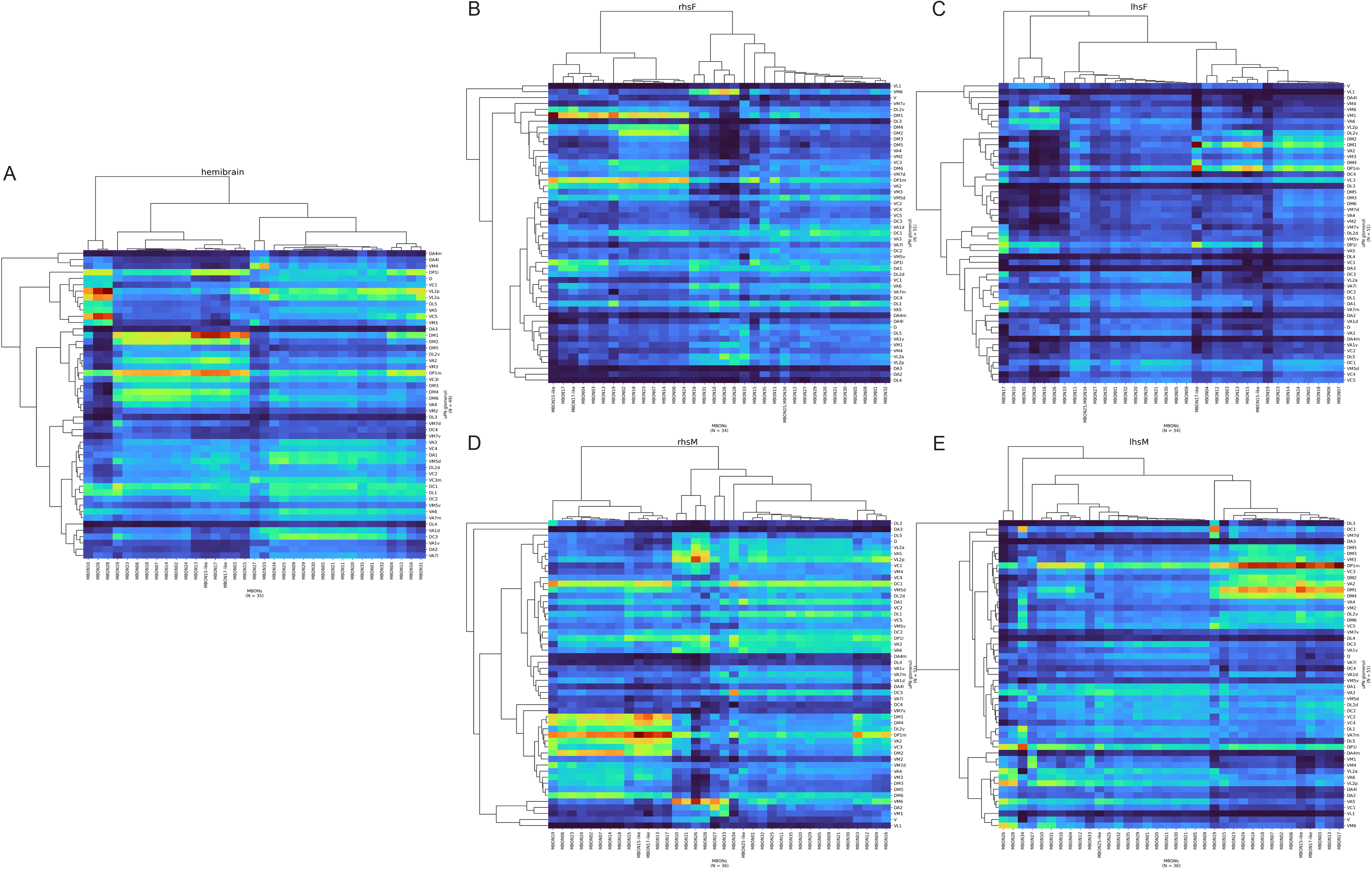
Cosine connectivity clustering of the effective connectivity from uPN glomeruli to MBONs in **(A)** hemibrain, **(B)** rhsF, **(C)** lhsF, **(D)** rhsM and **(E)** lhsM, using average linkage.

**Supplementary Figure 6.2:**
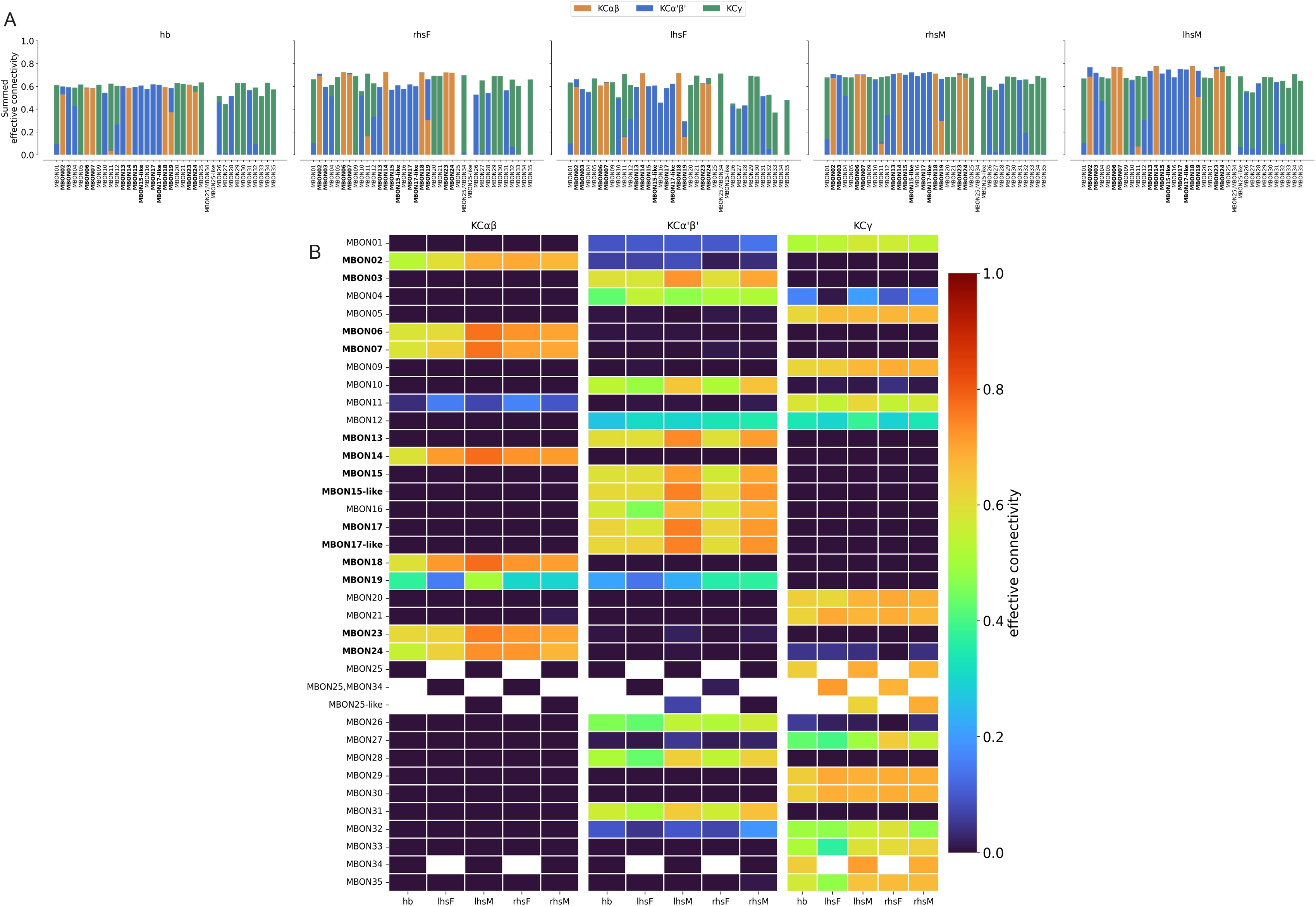
**(A)** KC-type composition of uPN-effective connectivity onto individual MBONs is consistent across five hemispheres. uPN→MBON effective connectivity was computed as the matrix product of normalised uPN→KC and KC→MBON connectivity matrices (uPN→KC normalised by each KC’s total Calyx input budget across all presynaptic partners, KC→MBON normalised by each MBON’s total KC-derived input; see Methods), then decomposed by KC type (KCαβ, KCα′β′, KCγ) by subsetting the KC→MBON matrix prior to multiplication. For each MBON, bars show the summed uPN-effective input attributable to each KC type (summed across all uPNs), stacked and coloured by KC type (KCαβ = orange, KCα′β′ = blue, KCγ = green). **(B)** Absolute magnitude of KC-type-specific uPN-effective connectivity onto MBONs across hemispheres. Heatmaps show summed uPN-effective connectivity from each KC type (KCαβ, KCα′β′, KCγ; left to right) onto each MBON (rows), across the five hemispheres analysed (columns). MBON labels in bold denote group 2 MBONs receiving strong effective input from food-associated PNs. Colour indicates summed effective connectivity on a shared scale across all panels (colour bar, right).

**Supplementary Figure 7.1:**
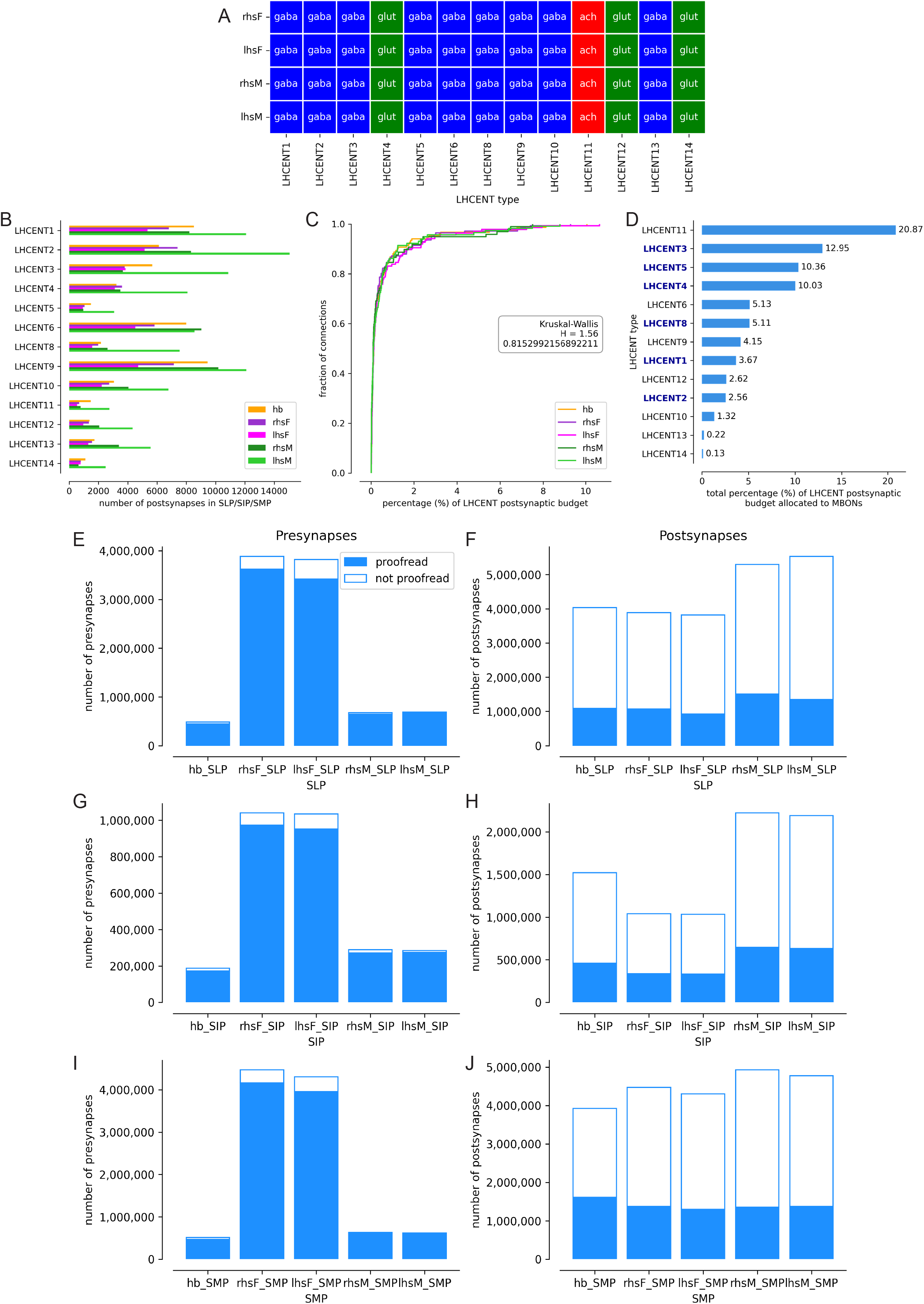
**(A)** Neurotransmitter predictions for each LHCENT type in FlyWire and mCNS. **(B)** Number of postsynapses each LHCENT type has in the superior protocerebral neuropils (SLP/SIP/SMP), across hemispheres. **(C)** Empirical cumulative distribution function (ECDF) plot of normalised MBON→LHCENT connectivity in the superior protocerebral areas, with a Kruskal-Wallis (KW) test. The KW test returned a test statistic of 1.56 and *p*-value of 0.815 (i.e. > 0.05). **(D)** Total proportion of input each LHCENT type receives from MBONs, using the averaged connectivity matrix in Figure 7B. LHCENT types in bold are those with recurrent projections back to the Calyx. Number of proofread and non-proofread presynapses (left) and postsynapses (right) in the SLP (**E** and **F**), SIP (**G** and **H**) and SMP (**I** and **J**).

**Supplementary Figure 7.2:**
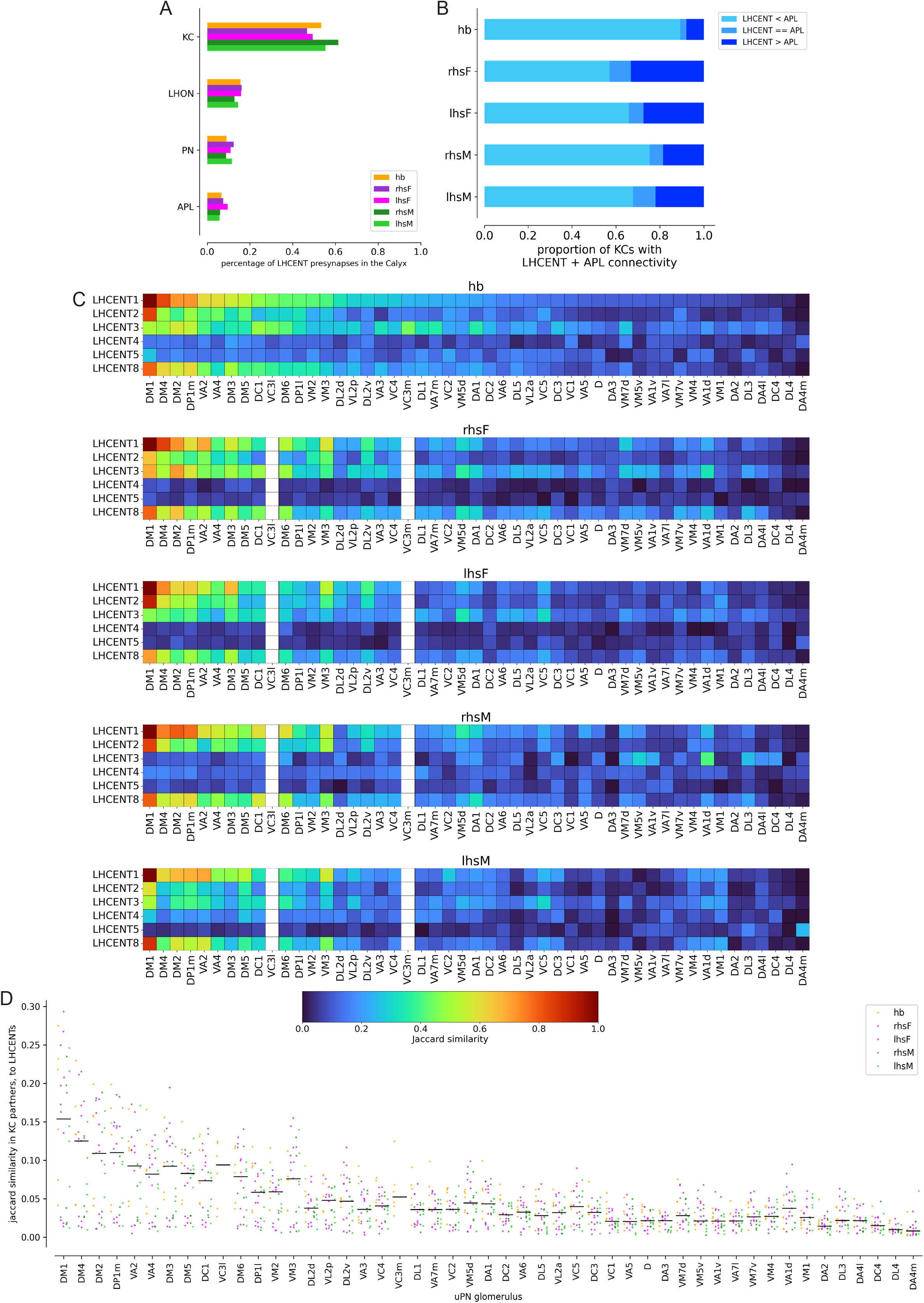
**(A)** The percentage of LHCENT presynapses onto different downstream cell types in the Calyx, across hemispheres. **(B)** The proportion of KCs for which total LHCENT input is less than, equal to or greater than APL input, across hemispheres. **(C)** Mean Jaccard similarity between LHCENTs and uPN glomeruli for the KCs they co-target in the Calyx, across hemispheres. **(D)** Distribution of Jaccard similarity values between uPN glomeruli and LHCENTs for the KCs they co-target in the Calyx; black vertical lines represent the means, presented in (C).

**Supplementary Table 1. Statistical comparison of odour-scene uPN input distributions.** For each KC, normalised connection weights from all uPNs assigned to a given odour scene were summed, with KCs receiving no input from that scene assigned a value of zero. Within each of the nine odour scenes and three KC types, per-KC percentage input distributions were compared across five hemispheres using Kruskal-Wallis tests, giving 27 omnibus comparisons; *p*-values were corrected across these tests using the Benjamini-Hochberg procedure, with (*q*<0.05) considered significant. Omnibus effect sizes are reported as epsilon-squared (∊^2^): negligible, <0.01; small, 0.01–<0.06; moderate, 0.06–<0.14; and large, ≥0.14. Significant omnibus tests were followed by pairwise Dunn’s tests between hemispheres with Holm correction; median differences and Cliff’s delta are reported as pairwise effect sizes.

## Acknowledgements

We would like to thank Catherine Whittle, Daniel Han and Davi Bock for comments on previous versions of the manuscript. We thank S. Waddell for discussions of the MB organisation. This study was supported by core funding from the MRC (MC-U105188491), Wellcome Trust collaborative awards 220343/Z/20/Z and 221300/Z/20/Z and a NeuroNex award (MRC MC_EX_MR/T046279/1). L.S.C. declares a financial interest in Aelysia Ltd.

## Author Contributions

All analyses were performed by M.W.P. who wrote the manuscript with input from G.S.X.E.J. and K.D.L. P.S. provided directions for some analyses. L.S. reconstructed and annotated KCs in the mCNS dataset; we thank the FlyEM Project Team and the Cambridge Connectomics Group for early access to these datasets.

